# Alignment of human behavior, brain, and AI models in the high-level valence processing of complex social scenes

**DOI:** 10.1101/2025.05.08.651182

**Authors:** Elahe’ Yargholi, Laurent Mertens, Laura Iris Van Hove, Ensa Mualla Gunay, Joost Vennekens, Jan Van den Stock, Hans Op de Beeck

**Affiliations:** Department of Brain and Cognition, Leuven Brain Institute, Faculty of Psychology & Educational Sciences KU Leuven, 3000 Leuven, Belgium; KU Leuven, De Nayer Campus, Dept. of Computer Science, J.-P. De Nayerlaan 5, 2860 Sint-Katelijne-Waver, Belgium; Leuven.AI - KU Leuven Institute for AI, 3000 Leuven, Belgium; Neuropsychiatry, Department of Neurosciences, Leuven Brain Institute, KU Leuven, 3000 Leuven, Belgium; York University, Department of Biology; Vrije Universiteit Brussel, Brussels, Belgium; Flanders Make@KU Leuven, 3000 Leuven, Belgium; Geriatric Psychiatry, University Psychiatric Center KU Leuven, 3000 Leuven, Belgium

## Abstract

Humans can evaluate the emotional meaning of complex social scenes in real-life settings. Recent evidence from the human brain and AI models pointed to visual processing as a core substrate for representing emotional valence of natural images, but this conclusion may not generalize to complex social scenes. We implemented experiments with social scenes in which emotional valence is partially dissociated from visual characteristics, objects, and scene settings. Human behavior, neuroimaging, and visual AI models confirm that visual processing captures basic emotional associations of objects and scene elements. However, when the valence of social scenes is incongruent with these basic properties, higher levels of processing are needed in the human association cortex and AI models. Our results show how and when valence processing demands advanced cognitive, neural, and computational processes that extend beyond the encoding of visual features.

## Introduction

Social cognition is crucial in human society, including the ability to read people’s emotions and intentions, experience empathy, and successfully interact with others and society at large. These capacities necessitate a reliable and fast understanding of the social and emotional meaning of events that happen around us. It has been shown that social and nonsocial emotional stimuli activate overlapping brain regions with additional cortical activation associated with mentalizing and prediction in social processing^1^. However, no comprehensive neural and computational studies have characterized how such emotional assessments can emerge from sensory input for highly complex social events.

In simple cases, emotion appraisals can be based on specific aspects of the environment. These cues can include non-social aspects, such as a general association of flowers and snakes with respectively positive and negative appraisals, or be of a more social nature, such as face or body expressions. A large literature has shown how these emotionally relevant cues are processed in the sensory cortex, for visual cues starting in the primary visual cortex and extending to object-, face-, and body-selective areas ^2–6^. Visual processing results in the categorization and identification of elements in a scene (e.g., is this a flower or a face? Is the face smiling?), and is further modulated by the emotions associated with these visual cues, with sometimes a higher visual activity for cues associated with negative emotions (negativity bias) ^7,8^ or positive emotions (pleasure bias) ^9^. Computational models such as Convolutional Neural Networks (CNN) trained on large databases of visual images and their associated emotions are able to generalize emotional inferences to new images^4,10,11^. In the case of natural scenes, these models are derived from more basic models trained to classify object images and one could hypothesize that they might strongly rely upon object-emotion associations (e.g., flower = ‘happy’).

However, for both the biological brain and AI models, visual processing might be insufficient for complex visual images that are of a social nature. For social events, we need to process a multitude of interactions between cues. In particular, while the literature has long been dominated by a focus on single cues such as the recognition of facial expressions, accumulating evidence suggests that for instance also body language and scene context are key elements to social cognition^12–15^.

In this study, we examine the limitations of visual processing in the brain and AI models to capture the behaviorally annotated valence of complex social scenes that require the processing of interactions between cues. We present a new experimental stimulus set with images that depict human social interactions in emotional scene contexts. Behavioral experiments show that across the full set, the valence of the people in the scene is partially dissociated from the valence of the scene context (Fig. 1). This is achieved by including incongruent images, e.g., people crying at a wedding. Next, we obtained neural responses using functional magnetic resonance imaging (fMRI) and examined the representation of emotional valence in the human brain as well as visual and multimodal AI models. FMRI data show that visual areas mostly represent visual properties and the emotional valence of the scene context, but not the valence of people in the scene or of the image as a whole. Neural signals related to the valence of the image as a whole are distributed across the association cortex, allowing the decoding of image valence and its generalization to new images even for incongruent images. Similarly, visual artificial neural networks trained to process valence rely mostly on the valence of the scene context while advanced multimodal AI models that integrate text and vision can partially capture the valence of the social interactions on top of the valence of the scene context. These findings reveal how understanding complex social interactions requires advanced cognitive, neural, and computational processes that go beyond the coding of visual features.

**Fig. 1.**
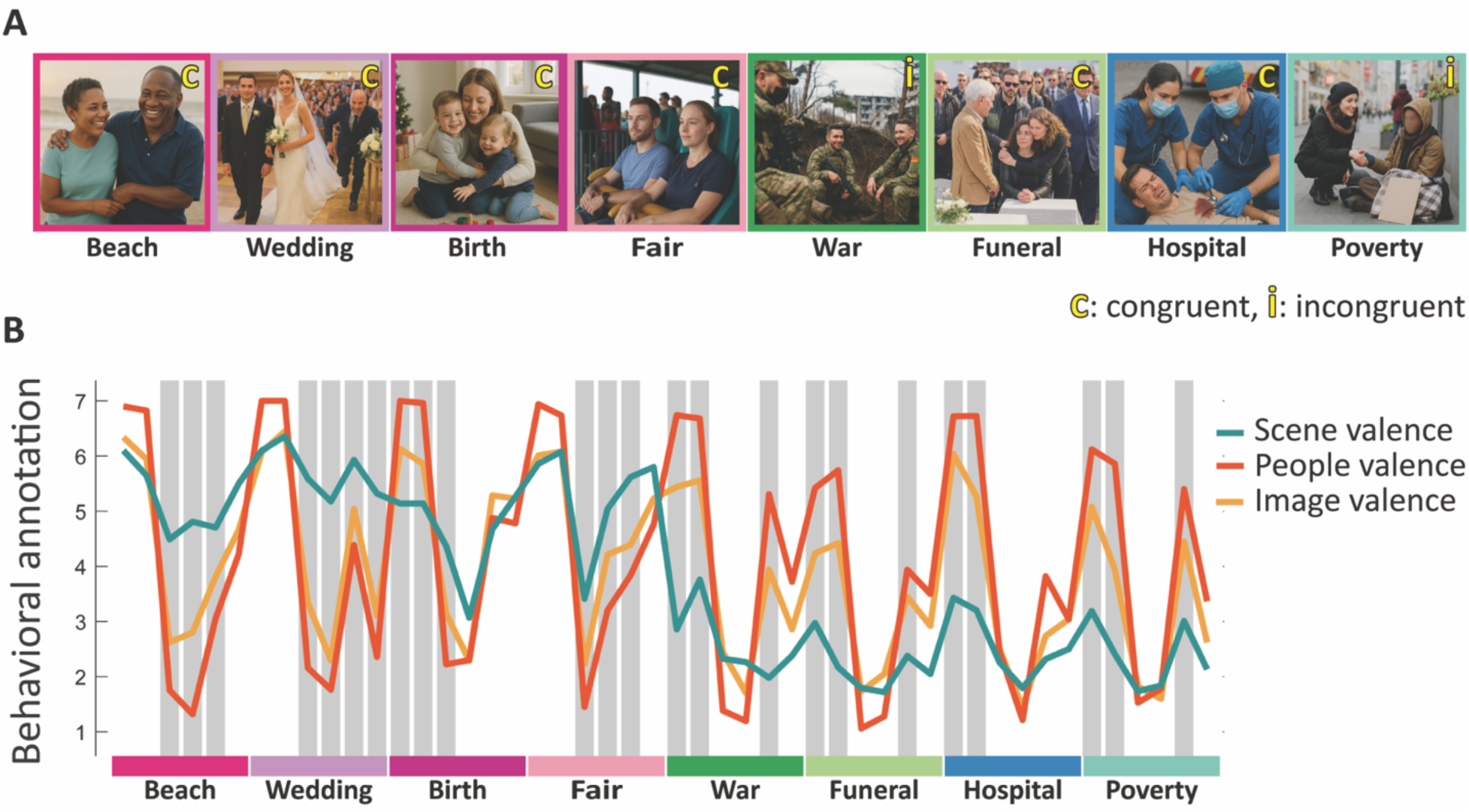
Stimulus design and behavioral annotations. A, 8 stimulus images out of the set of 48 stimuli (original images are replace with a computer generated images to have people unidentifiable), color-coded based on the valence of the scene context and labeled (i)c for (in)congruent valences of the scene and people depicted in the scene. Links to all original images are provided in Suppl. Table S1. B, Behavioral annotations for all stimuli. Estimated reliability [with a confidence interval of 95%] for scene, people, and image valence was respectively 0.99 [0.98,0.99], 0.99 [0.98,0.99], and 0.99 [0.98,0.99]. The gray background indicates stimuli with incongruent valences of the scene and people depicted in the scene.

## Results

### Social scene images that dissociate image valence from the valence of the scene context

We compiled a large set of images that depict social interactions in a variety of emotionally relevant settings. Based on behavioral annotations, we created a final stimulus set that comprised 48 complex social scenes (Fig. 1). We obtained valence annotations for the scene (background, objects, and clothing), people depicted in the scene (facial and body expression), and the image as a whole in 50 participants (Fig. 1B). The first half of images include positive scene contexts (e.g., a wedding), while the second half of images convey a negative scene context (e.g., a funeral). For some images, referred to as congruent images, the valence of the scene context is very similar to the people and image valence. For other, incongruent images (grey bars in Fig. 1B), scene valence and people valence are very different. For these incongruent images, image valence always ends up in between scene and people valence.

### Cross-stimulus decoding reveals the limitation of low-level ROIs in predicting valences

To evaluate the representation of scene/people/image valence in different brain regions, we employed a decoding approach. Using ridge regression, we fitted the decoding models to the fMRI data acquired while subjects viewed the stimulus images and rated the valence of the whole image. We predicted positive vs. negative valences for scene/people/image in HCP (Human Connectome Project) regions following a cross-stimulus leave-one-run-out approach (Fig. 2A). We adopted a cross-stimulus decoding approach, i.e. we used subsets of stimuli for training the decoder and other stimuli for testing the decoder prediction. Decoding performance was evaluated by calculating the Pearson correlation between true and predicted valences ^16^ and tested for significance using one-sample one-sided t-tests corrected for multiple comparisons across ROIs (FDR) ^17^. We summarized the results considering a division of HCP regions into three large regions of interest (ROIs) (Fig. 2B): (i) low-level visual regions (retinotopic areas V1-V4), (ii) mid-level association areas mostly in posterior cortex, and (iii) high-level ROIs involved in control processes and mostly in anterior cortex. We averaged decoding performances across their constituting HCP regions. Statistical tests, one-sample one-sided t-tests corrected for multiple comparisons across ROIs (FDR) (Fig. 2C), showed a significant correlation between true and predicted labels for all the valence sources in mid/high-level regions, but only for scene valence in low-level regions. The variation of cross-stimulus decoding across source of valence and ROIs was further tested in a 2-way ANOVA with source of Valence (scene/people/image) and Region (low/mid/high-level) as within-subject factors. We found a significant main effect of Valence (F(2, 84) = 3.96, p = 0.03) and a significant Valence*Region interaction (F(2, 84) = 3.60, p = 0.01), but not of region (F(2, 84) = 0.40, p = 0.67). The interaction reflects that low-level visual areas represent the emotional valence of the scene context and not the valence of people in the scene or the valence of the whole image which are represented only in higher-level areas.

**Fig. 2.**
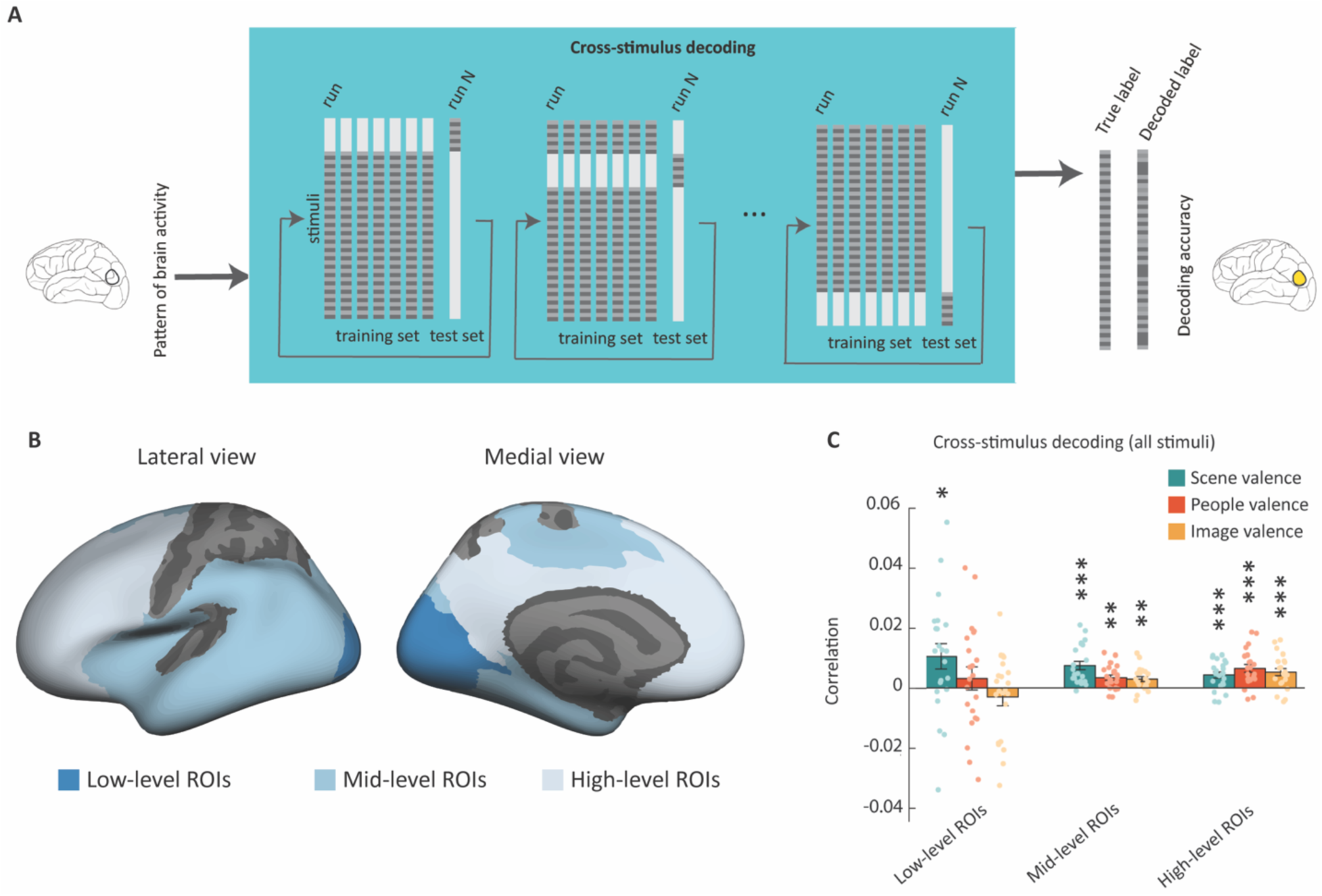
Cross-stimulus decoding approach and performance. A, Schematic presentation of cross-stimulus decoding approaches. Decoders predicted positive vs. negative valences following a leave-one-run-out approach. In cross-stimulus decoding, subsets of stimuli were used for training the decoder and other stimuli for testing the decoder predictions. B, Division of HCP ROIs into three large ROIs (see Methods for more details). C, Performance of cross-stimulus decoding for scene/people/image valence in low/mid/high-level ROIs. The bar plots show the mean decoding accuracy (Pearson correlation between true and predicted valences). Dots represent individual subjects and error bars indicate SEM (* p < 0.05, ** p < 0.01, *** p < 0.001, one-sample one-sided t-test, FDR-corrected).

To further pinpoint the location of the strongest cross-stimulus decoding results, we applied a previously proposed clustering of HCP ROIs ^18^ in 19 regions covering the entire cortex except the somatosensory and motor, early auditory, and superior parietal regions. Scene valence was significantly predicted across stimuli in posterior ROIs (ventral stream visual, insular and frontal opercular, temporo-parieto-occipital junction, inferior parietal) (Suppl. Fig. S1). The posterior cingulate significantly decoded both scene and people valence. The only other significant prediction for people valence was derived in a more anterior ROI, dorsolateral prefrontal. No region showed significant decoding for image valence. Given the highly significant decoding for image valence when pooling predictions across smaller ROIs into a large mid-level and high-level ROI, the neural signals related to image valence seem to be distributed across a larger set of regions.

The decoding across stimuli is important to rule out the possibility that the results might be dominated by strong effects of subsets of stimuli and visual features that tend to correlate strongly with valence. Indeed, simple decoding (Suppl. Fig. S2A) shows a significant correlation between true and predicted labels in low/mid/high-level regions for all valences, with higher correlations for low-level ROIs (primary visual and early visual) and scene valence (Suppl. S2B & 2C), not due to low-level features (Suppl. Fig. S3).

The inclusion of incongruent images is also crucial to show the limitations of low-level processing as a basis for valence assessment. We applied cross-stimulus decoding to congruent and incongruent stimuli separately (Fig. 3A). For congruent stimuli, we observed significant correlations between true and predicted labels for all the valences in low/mid/high-level regions, with stronger correlations in low-level ROIs. This finding mimics the results for simple decoding across all stimuli. In contrast, for the incongruent stimuli, we did not observe significant correlations in low-level visual regions, not even for scene valence. In the mid- and high-level ROIs, the representation of valence seems sufficiently abstract to generalize across images, both for congruent and incongruent images.

**Fig. 3.**
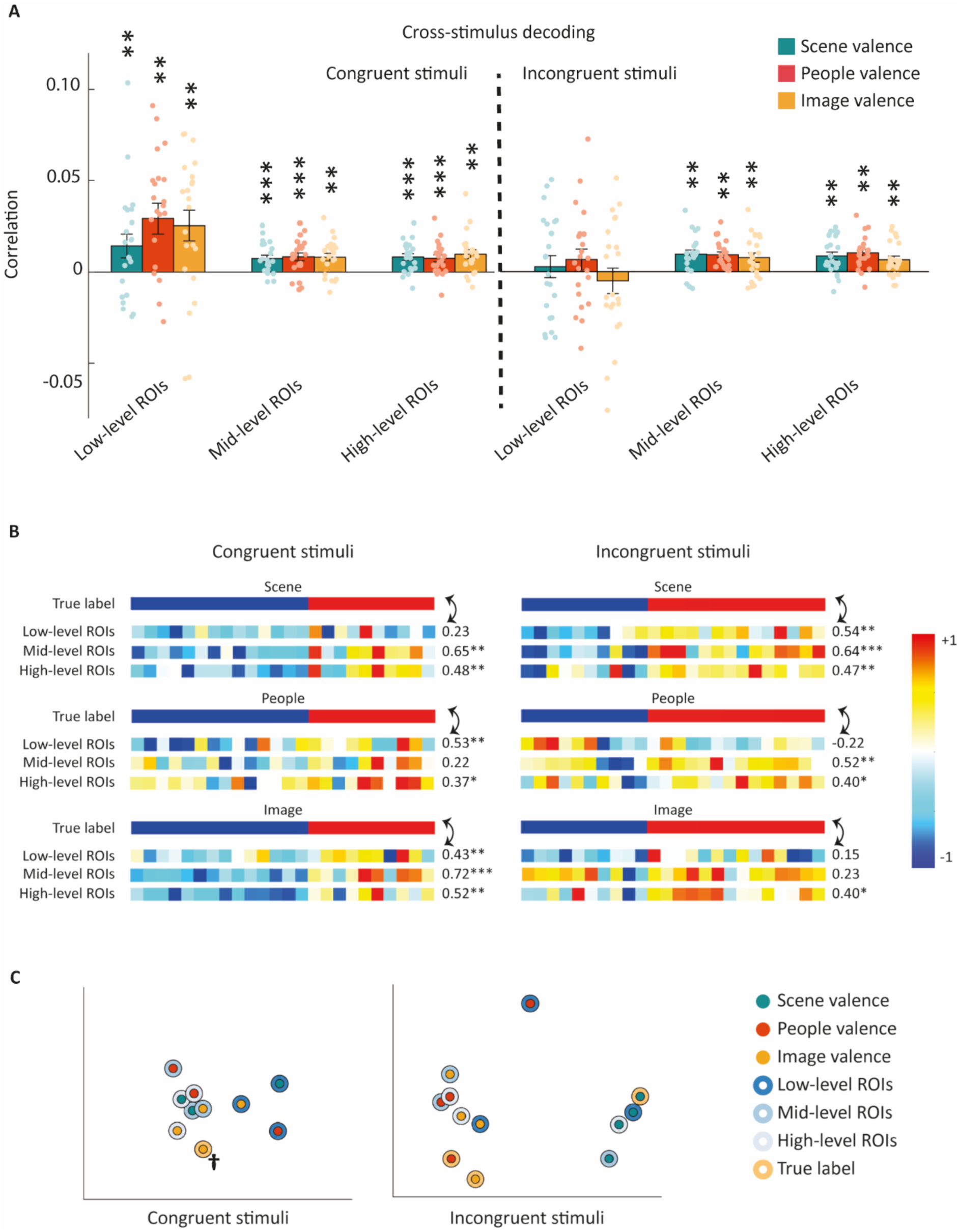
Performance of cross-stimulus decoding in low/mid/high-level ROIs for congruent and incongruent images separately. A, Performance of cross-stimulus decoding. The bar plot shows the mean decoding accuracy (Pearson correlation between true and predicted valences) in ROIs for scene/people/image valences. Dots represent individual subjects and error bars indicate SEM (** p < 0.01, *** p < 0.001, one-sample one-sided t-test, FDR-corrected). B, Comparison (correlations) of behaviorally annotated valences (“true label”) with brain-decoded scene/people/image valence from cross-stimulus decoding, now averaging across participants prior to the correlation with behavior (“group-averaged” predicted labels) (* p < 0.05, ** p < 0.01, *** p < 0.001, one-sample one-sided permutation test). Each square represents an image and they are sorted to have all negative true labels at the left side and all positive true labels at the right side. C, Two-dimensional similarity structure as obtained by applying MDS to the distance matrix of the true label and brain-decoded valences from cross-stimulus decoding shown in panel B. Points are color-coded based on valence source (inner circle for scene/people/image valence) and the true/brain-decoded labels (outer circle for true label or brain-decoded predictions from low/mid/high-level ROIs). **†**For congruent stimuli the three points for the true label for scene/people/image valence fully overlap.

### Large effect size when predicting behavioral valence ratings from neural representations

Inspired by the significant cross-stimulus decoding in mid- and high-level regions for complex social scenes, we wondered how closely we could predict behavioral ratings of valence for these images from the brain data. In previous analyses, the decoding represents data at the individual image level and the individual participant level. If the neural representations underlying the cross-stimulus decoding capture variation in behaviorally rated valence in a meaningful way, then we would expect that decoding performance would go up markedly when predictions are averaged across participants to increase reliability. We averaged the predictions from the cross-stimulus decoding across participants (the “group-averaged” predicted labels) prior to correlating the predicted labels with the behavioral ratings (which we refer to as the “true labels”).

We evaluated the reliability of the group-averaged predicted labels by computing the split-half reliability. We randomly divided participants into two groups, averaged predictions across participants in each group, computed the correlation between them and applied Spearman-Brown correction. Estimated reliability for mid-level and high-level ROIs, averaged across all sources of valence, was respectively 0.47 and 0.45, compared to an average reliability for low-level ROIs of 0.19.

If the neural representations capture a meaningful proportion of the variation in valence ratings across stimuli, then there should be a high correlation between the group-averaged predicted labels and the true labels, possibly up to the level of these reliability estimates. Figure 3B shows these correlations for the three aggregated regions used before. For congruent stimuli, predicted labels and true labels for scene/people/image valence were significantly correlated in most ROIs. For incongruent stimuli, we can observe a progressive change from low-level to high-level ROIs: significant correlations between predicted and true labels were present only for scene valence in low-level ROIs, while they are typically significant for all sources of valences in high-level ROIs. The correlations between the group-averaged predicted labels and the true labels in mid- and high-level ROIs were 0.47 on average, which is very close to the estimated reliability mentioned above. This outcome suggests that the true labels capture close to all variability in the fMRI predictions that is replicable across participants.

To further examine the similarity among predictions, we computed the correlation distances (one minus the correlations) and applied multi-dimensional scaling (MDS) analysis on the distance matrix of the vectors shown in Figure 3B. Figure 3C depicts the first two dimensions that capture most of the variance. The results confirm the similarity of predicted labels and true labels for scene/people/image valence for congruent stimuli. For incongruent stimuli, predictions for scene valence in low/mid/high-level ROIs were close to the true labels for scene valence and formed a separate cluster away from the other vectors related to people and image valence.

These patterns confirm the conclusions from the decoding at the individual level in previous analyses, with decoding of people and image valence in mid- and high-level regions, even when dissociated from scene valence. In addition, the analyses on group-averaged predicted labels reveal that the predicted labels correlate highly with true labels when predictions are averaged across participants.

### Category-selective regions are more sensitive to the visual features of their favorite categories than valences

Faces, bodies, and scenes are obviously highly relevant for processing the images in our stimulus set, as these categories are present in every image and carry relevant information concerning emotional appraisals. Part of the mid-level ROIs includes well-known visual areas with selectivity for these visual categories ^19^, as well as areas covering the superior temporal sulcus recently suggested to specifically process human interactions ^20,21^. Could we be inadvertently disregarding a potential co-localization of valence representations with neural selectivity for these highly relevant stimulus categories? To examine the role of category-selective regions in emotional assessment, localizer data (recorded in the same subjects) and anatomical/functional masks were employed to determine V1, category-selective networks for faces, bodies, and scenes, and the superior temporal sulcus. We computed Spearman’s correlations between univariate responses in these regions and visual features related to depicted faces and bodies in the images (e.g., size/number of faces and bodies, Fig. 4A). As expected, univariate responses of category-selective networks correlated with the visual features of their favorite category (Fig. 4B). Univariate responses in the body network were significantly correlated with the total image area of all depicted bodies relative to the image size (Body-Area) and not correlated with the number of depicted bodies (Body-Count). Univariate responses in the face network were significantly correlated with the total area of all depicted faces relative to the image size (Face-Area, Fig. 4B & 4C) and negatively correlated with the number of depicted faces (Face-Count) and the average distance between face pairs relative to the image size (Face-Distance). Univariate responses in the scene network showed the opposite profile to those of body and face networks (Fig. 4B & 4C): a negative significant correlation with face/body-area and a positive significant correlation with face/body-count and face-distance. Thus, despite the fact that all the images in our stimulus set contain faces, bodies, and scene background, there is sufficient variation in the properties of these image elements to result in strong variations among images in terms of how much they activate the different category-selective regions in a way that fits with what one could expect given the preferred category of each region.

**Fig. 4.**
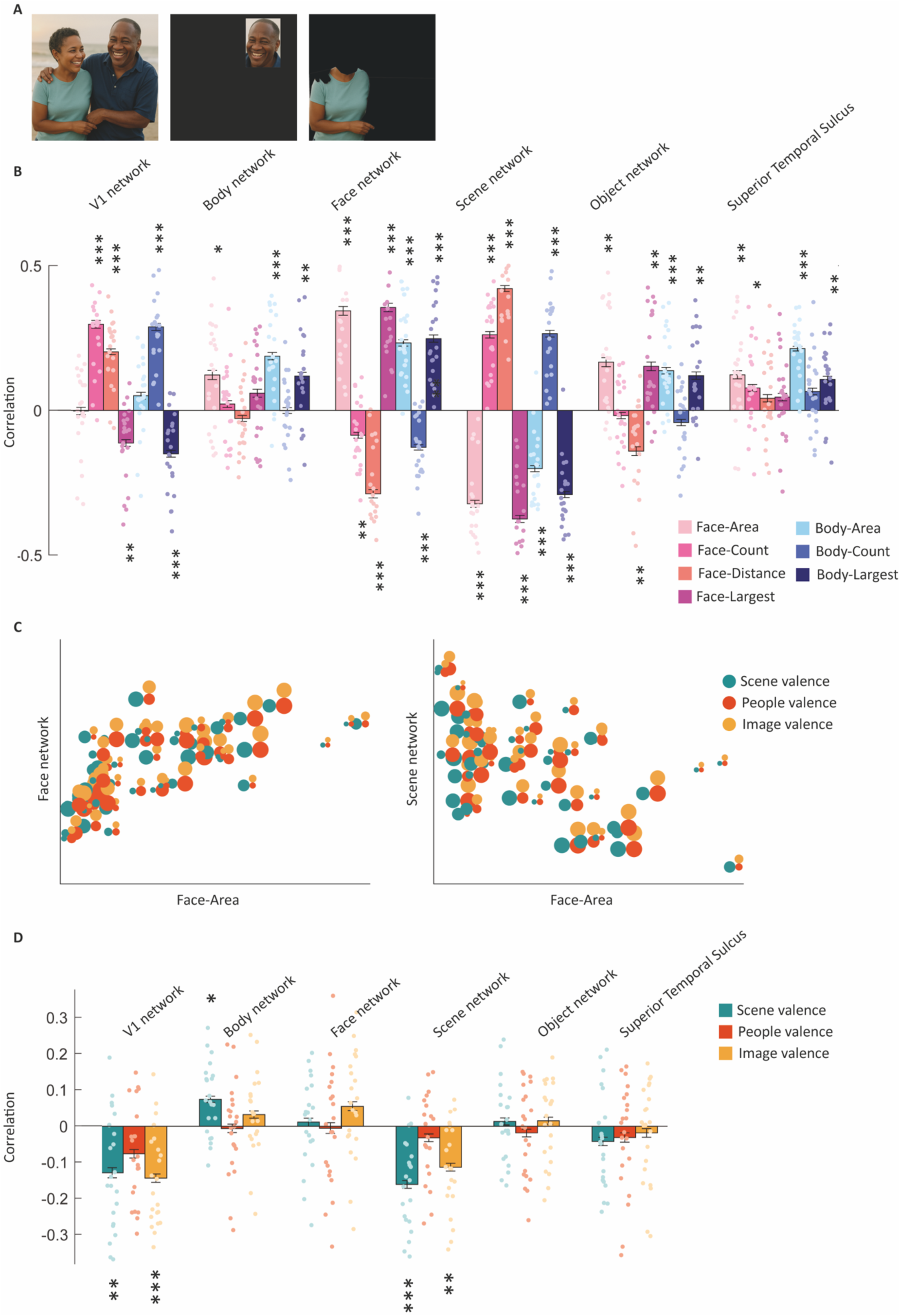
Role of category-selective regions in representing the properties of complex social scenes. A, An example of a face and body extracted for a stimulus image. Extracted faces and bodies were used to quantify visual features like the size or number of depicted faces and bodies for our stimuli (original image is replace with a computer generated image to have people unidentifiable). B, Correlation between univariate responses of category-selective networks and visual features. Bar plots represent the mean Spearman’s correlations between univariate responses of category-selective networks and visual features. Dots represent individual subjects and error bars indicate SEM (* p < 0.05, ** p < 0.01, *** p < 0.001, one-sample one-sided t-test, FDR-corrected). C, Univariate responses in the face and scene network. Univariate responses in the face network were significantly correlated with the total area of all depicted faces relative to the image size (face-Area) while univariate responses in the scene network showed the opposite profile. Each stimulus image is presented with three connected circles. The color of the circles represents valence source (scene/people/image valence) and the size of the circle shows the quantity of corresponding behavioral annotations. D, Correlation between univariate responses of category-selective networks and annotated valences. The bar plot represents the mean Spearman’s correlations between univariate responses of category-selective networks and annotated valences. Dots represent individual subjects and error bars indicate SEM (* p < 0.05, ** p < 0.01, *** p < 0.001, one-sample one-sided t-test, FDR-corrected).

These strong variations in neural response as a function of stimulus properties contrast in effect size with the much smaller variations found for valence. Following a similar approach, we computed the correlations between univariate responses and behaviorally annotated valences (for scene/people/image). Results (Fig. 4D) showed significant negative correlations with scene and image valence in V1 and the scene network, revealing that negative scene and image valences give higher activity. Note that this negativity bias was not confounded by visual features, as there were no significant correlations between valence and the visual features (for scene valence even not without FDR correction; for image valence there were moderate correlations with Face_count, r=-0.27, p_uncorrected_ = 0.040, and Body_Count, r=-0.28, p_uncorrected_ = 0.029). Most importantly, the size of the correlations with valence in Figure 4D is much smaller than the correlations with visual features in Figure 4B. In particular, the absence of any correlations with people valence in body- and face-preferring networks illustrates the extent to which these regions represent visual features of faces and bodies rather than their emotional valence when presented in a complex natural stimulus.

Finally, we also employed cross-stimulus decoding to predict positive vs negative valences for scene/people/image in these regions. Results did not reveal any significant cross-stimulus decoding, except for scene valence in the scene network (Suppl. Fig. S4). These data showed that emotion perception in category-selective networks was limited to the effect of scene valence in the scene network. Valence predictions are more robust when aggregated across larger cortical territory (see previous section and Fig. 3), and regions that are selective for faces and bodies do not seem to be hotspots for decoding the valence of instances from these categories when presented in a complex natural scene.

### How do AI models represent the valence of complex social scenes?

We compared the reported behavioral and neural representations of our stimuli with their representation in AI models. We examined representations in the last hidden layers, verbal descriptions, and valence annotations/space across various AI models including simple visual models like AlexNet ^22^ and EmoNet ^4^, and more advanced multimodal models such as CLIP ^23^ and Large Language Models like Llama ^24^ and GPT4 ^25^. In literature, EmoNet was suggested to be a benchmark model for capturing valence in natural images. The visual processing and training of CLIP is still comparable with AlexNet and EmoNet (convolutional architecture; trained with images of similar complexity), yet it is a more modern model that also received verbal labels of these images and shows a higher correspondence with neural representations ^26^. Llama is an open-source multimodal large language model with an extensive training set and a large number of parameters. We include GPT4 in some analyses, despite not being an open model, because it is arguably the state of the art and hence a failure or success of this model is a useful benchmark.

#### Alignment between behaviorally evaluated valence and image valence predicted from AI models

We used a similar approach as implemented for the human fMRI data by extracting valence predictions from the models and comparing these valence predictions with the behavioral annotations. The extraction of the valence predictions was adjusted to the characteristics of each AI model. First, for EmoNet, Kragel et al., 2019 constructed a predictive model of image valence from the last fully connected layer of EmoNet using partial least square regression. Second, for CLIP, we augmented the model with a single layer at the output and finetuned it on the large FindingEmo dataset containing complex social scenes to provide a prediction for image valence (for more details see Method). Third, we prompted Llama and GPT4 for scene/people/image valence ratings, giving the models the same instructions (different prompts for different sources of valence) and definitions for valences as provided to human annotators (data included for 40 images in correlations with GPT4, as GPT4 refused to rate valences for 8 images). Then, we computed the (Spearman) correlation across images between valence predictions from AI models and behavioral annotations for scene/people/image valence.

For congruent images, all behavioral annotations and AI valence predictions were correlated (left side of Fig. 5A&C). A few examples scatter plots are visualized in Fig. 5C-left. Incongruent images provide a unique test, as behaviorally annotated scene and people valence for these images are not positively correlated (even slightly negative), nor are scene and image valence (see top lines in Fig. 5A-right). Investigating these incongruent stimuli reveals differences between AI models in their understanding of the different sources of valence. All models showed a positive correlation with behavioral scene valence (except when GPT4 was prompted to provide people valence). Image and people valence were more difficult to capture. Image valence predicted by EmoNet and behavioral annotation of image and people were not positively correlated, with even a slight negative correlation (Fig. 5B left). In contrast, predictions from CLIP and ratings of Llama and GPT4 were almost equally correlated with behavioral ratings for image valence (Fig. 5A-B). Also, for capturing people valence, EmoNet was an outlier in the negative direction, given that its prediction negatively correlated with behavioral people valence, while Llama and GPT4-People showed a positive correlation with behavioral people valence (Fig. 5B). CLIP-image also tended to show a positive correlation with behavioral people valence. When compared directly, CLIP-image, Llama-image and GPT4-Image did not differ in their correlation with behavioral people valence in incongruent images (permutation test, p > 0.100), while all showed a significantly higher correlation with behavioral people valence compared to EmoNet-Image (permutation test, p < 0.010) (Fig. 5C).

**Fig. 5.**
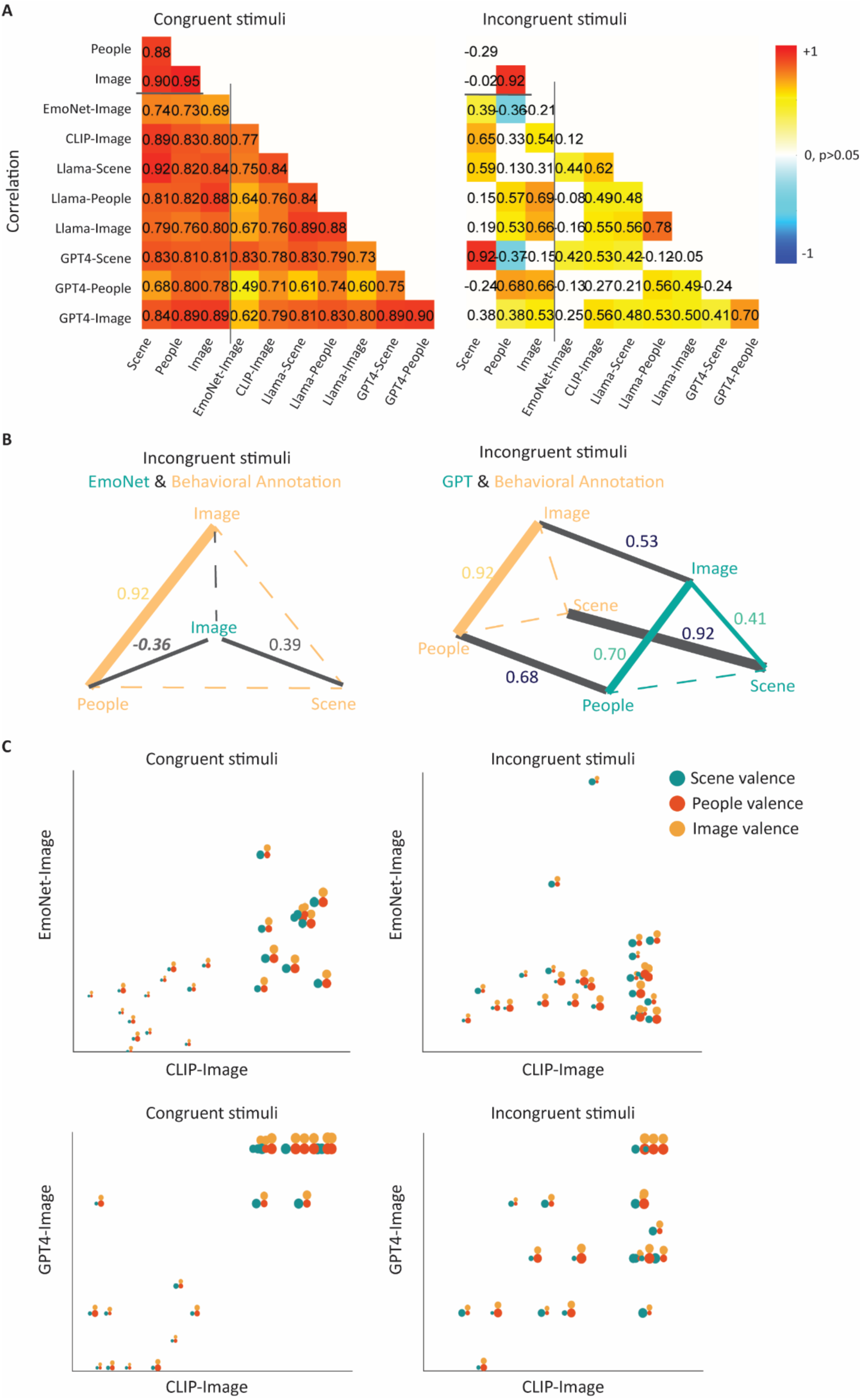
Comparison of behaviorally evaluated valence and valence predicted from AI models. A, Correspondence between behavioral annotations and AI predictions for valence. Colored cells in the correlation matrices are significant (p < 0.05, permutation test). Values above the horizontal black line are correlations merely among behavioral RDMs and values on the right of the vertical black line are correlations merely among model RDMs. B, Comparison of behavioral annotations and AI predictions for valences of incongruent stimuli, for a subset of comparisons. Values and line thicknesses represent the size of correlations between the valences for EmoNet-Image (left) and GPT4 (right) (p < 0.05, permutation test), with dashed lines referring to insignificant correlations. C, Scatter plots showing the predicted valence for individual images of EmoNet-image, CLIP-image and GPT4-Image. Each stimulus image is presented with three connected circles. The color of the circles represents valence source (scene/people/image valence) and the size of the circle is proportional to how positive the corresponding behavioral annotations are (large = positive).

#### Alignment between neurally predicted valence and AI-predicted valence

After these comparisons between AI models and behavior, we additionally compared the valence predictions of AI models with the group-averaged valence predictions based upon multivariate neural responses in low/mid/high-level ROIs (Fig. 6). For congruent stimuli, all models (EmoNet, CLIP, Llama and GPT4) provide predictions significantly correlated with neurally predicted valences (Fig. 6A-left) and these correlations increase with higher levels of model complexity (smallest for EmoNet).

**Fig. 6.**
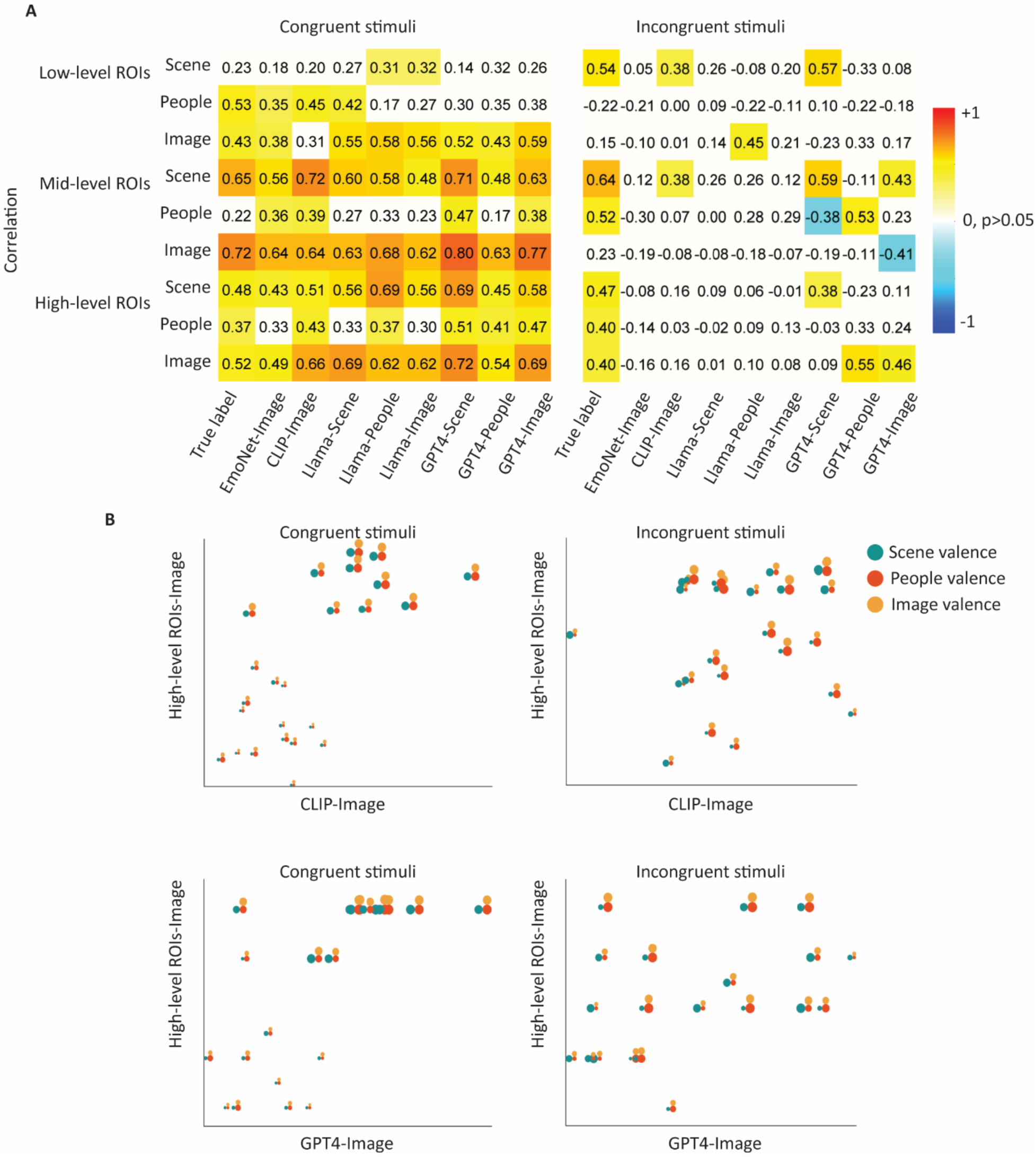
Comparison (correlations) of AI-predicted and behaviorally annotated valences (“true label”) with brain-decoded scene/people/image valence from cross-stimulus decoding in low/mid/high-level ROIs. A, the Correlation matrix of the valences. Colors represent significance, with colored cells passing the threshold of p < 0.05 (permutation test). The first column duplicates the results shown before in Fig. 3B (correlations between brain & behavior), the other columns directly compare brain and AI predictions. B, Representation of brain-decoded image valence from cross-stimulus decoding in high-level ROIs vs. CLIP/GPT4 prediction for image valence. Each stimulus image is presented with three connected circles. The color of the circles represents valence source (scene/people/image valence) and the size of the circle shows the quantity of corresponding behavioral annotations. *The role of explicit prediction of valence: Overall similarity relationships in human behavior and AI models fail to capture people and image valence in the presence of incongruence*

For incongruent stimuli, the EmoNet prediction was not significantly correlated with neurally predicted valence in any of the ROIs and for any of the sources of valence (Fig. 6A-right). CLIP-image predictions were correlated with neural decoding for scene valence in low/mid-level ROIs but failed to capture neural predictions for people and image valence (Fig. 6, A and B). Thus, while neural and CLIP-Image predictions both related to behaviorally annotated image valence in the previous section, CLIP-Image fails to align with the neural representation of image valence for incongruent stimuli. With both Llama and GPT4, we could employ prompt engineering, but we see a clear contrast between the two language models (Fig. 6B). Only GPT4 ratings result in correlations with all sources of valence as predicted neurally. GPT4-Scene correlates with neural predictions of scene valence in all ROIs. Correlations of GPT4-People and GPT4-Image vary across ROIs, with for example significant correlations with neurally predicted people and image valence in the high-level ROI. In sum, only the largest multimodal language-vision model combined with prompt engineering succeeded in emulating neural decoding of valences. Yet, even for this model the success of the predictions was variable, with for example a significant negative correlation between GPT4-Image and image valence in mid-level ROIs.

We described how neural and AI representations incorporate people valence, allowing them to make sense of the valence of the total scene even for incongruent images where scene and people valence do not agree. These findings were obtained by explicitly probing for valence representations. In human behavior and for Llama or GPT4, we explicitly asked about the different sources of valence. In neural data and for AI models like EmoNet and CLIP, we trained classifiers to explicitly predict image valence. In contrast, when we use methods that capture overall representational similarity, without explicitly referring to properties like people or image valence, we find that only scene valence captures part of the similarity relationships. More details are presented in Suppl. note 6.

## Discussion

We investigated the cognitive, neural, and computational processes that underlie the ability to interpret complex social scenes in terms of valence. We used an experimental paradigm with social scenes that dissociate emotional valence from visual characteristics, objects, and scene settings. We obtained behavioral annotations and recorded responses from the human brain and AI models. Employing cross-stimulus decoders, we found that low-level visual areas represent just the scene valence while mid/high-level ROIs decode the people/image valence in addition to the scene valence. While the information regarding valence is present in the faces, bodies, and scene elements in the images, neural responses (fMRI) of regions selective for these categories show strong correlations with visual properties of these categories in the images but not the people and image valence. Comparison of the neurally predicted labels with true labels for incongruent stimuli reveals that in low-level ROIs, there is a significant correlation between predicted and true labels only for scene valence, whereas in high-level ROIs, all sources of valences show significant correlations. These findings show the superiority of mid/high-level over low-level ROIs in the assessment of the emotional valence of different components in a complex scene and their integration into an overall valence. Incongruent stimuli also challenged AI models. While all models align with the behavioral ratings of scene valence, image and people valence are only predicted by models that are not purely visual (CLIP, Llama and GPT-4). Neural predictions of image and people valence were even more very difficult to capture, with only the largest model (GPT4) having some but still variable success.

A large literature on visual perception of valence has shown that emotional valence of many types of images is represented as early as the primary visual cortex ^2–4,27,28^ and face-, body- and object-selective regions ^6^. However, these studies have mostly used stimuli like expressive human faces or random natural scenes. For these kinds of stimuli, emotional aspects correlate with low-level visual features or objects. Various studies have explicitly used such low-level information to estimate emotion dimensions or emotion categories^29–33^. These correlations could be the main cause of positive results for valence representation in visual areas. As a case in point, Kragel et al. ^4^ suggested that EmoNet (based on AlexNet) captures the emotional content of visual stimuli consistently with behavioral annotations. They also provided evidence on the representation of visual emotion schemas within the occipital lobe and partially redundant coding of this information in other brain regions. However, their different emotion schema was correlated with object classes of visual stimuli (classification of emotion categories based on object classes resulted in top 1 accuracy of 11.4% with chance level at 5%). Most recently, Abdel-Ghaffar et al., 2024 showed tuning of occipito-temporal activity to semantic and affective features, with an added value of affective features on top of the semantic model, and in combination these models were better than a visual Gabor model. However, their semantic model is very simplistic (presence of a small set of categories), as is the Gabor model, and they did not investigate mid-level structural features such as color or shape so many left-out visual attributes could correlate with the dimensions of arousal and valence. This is a typical problem with non-curated stimulus sets containing natural images. Specific experiments are needed to really understand how dimensions are computed and interact with each other^34^.

Here we presented such a paradigm. By manipulating scene context, we include a factor that covers the sort of associations that might underlie many of the findings in the literature. For example, a wedding dress is typically associated with positive emotions, in contrast to military gear. Given such associations, and all the findings in the literature that might rely upon these associations, it is not surprising that we can predict the valence of scene context from neural activity in the visual cortex. However, the valence of the full image can only be predicted from this visual activity for congruent images where all sources of valence go together. This is probably the situation in almost all images in the non-curated stimulus sets that have been used in prior studies. Specifically for natural stimuli with a social content and interactions, ^21,35^ showed that visual processing serves a key role in social-affective cognition, although by itself it may not be entirely sufficient. This result is consistent with our findings for congruent social scenes, but their stimulus sets did not include incongruent social scenes.

Our stimulus set includes incongruent images, in which we deliberately dissociate scene valence from image and people valence. For these images, multivariate activity in early visual regions does not help to predict image valence. Nevertheless, there is a distributed neural representation of image and people valence in mid- and higher-level ROIs. Similarly, the incongruent images have proven essential to test computational models. Models like EmoNet that have been developed to capture image valence with simpler natural scenes fail completely with more complex social scenes.

We observed in supplemental analyses (see suppl. Note 6) that training multivariate classifiers or specific prompt engineering is needed to uncover these valence representations. Methods that simply assess representational similarity (e.g., representational similarity analysis or GUSE semantic mapping) reveal representations that are strongly or sometimes even exclusively influenced by scene valence. The flexibility of multivariate decoding in reweighting individual features to predict a target variable and explicit and direct access provided by prompt engineering, in comparison with the “equal weight” assumption of representational similarity analysis (RSA), could be the reason for RSA inefficiency in this application ^36^.

We were surprised to find that neural predictions about image valence were more difficult to capture with AI models than behavioral annotations. The CLIP and Llama models correlated nicely with behaviorally assessed image valence but did not correlate with neurally predicted image valence, and even GPT4 had variable success. This is not the first observation in the literature that neural findings can be more difficult to model than human behavior. Recently, Gokce & Schrimpf, 2024, tested over 600 models trained in core object recognition and reported that behavioral alignment continues to scale with larger models, while neural alignment saturates. For the largest models, the behavioral alignment is better than the neural alignment. Further research is needed to understand these discrepancies, not only at the modeling level, but also at the level of cognitive neuroscience to understand what is different between neural representations and behavior.

We do not claim that we provide a full description of the perception of social scenes and their valence. In particular, we have ignored the subjective nature of such emotional assessments. First, different observers might differ in their assessment of emotional valence, behaviorally as well as neurally^38^. In our analyses, we focus on the aspects of these assessments that are common across observers, and we explicitly average behavioral findings and neural predictions across observers. Second, to overcome the noise inherent to fMRI measurements our decoding approach is based on the repeated presentation of stimuli and decoding accuracy depends on the consistency of responses across multiple representations. However, responses to stimuli in general and emotional stimuli in particular change with repeated exposure even within one and the same individual^32,39^. Therefore, our approach would be more successful for regions with less sensitivity to stimulus novelty across presentations (have a stationary BOLD response) and might underestimate the strength of the valence representations in regions sensitive to the effects of stimulus repetition.

## Conclusion

We investigated mechanisms underlying human evaluation of emotional meaning in complex social interactions. While visual processing plays a fundamental role in representing the emotional valence of simple natural images, evidenced by both human neuroimaging and AI models, it is incompetent when applied to complex social scenes. Advanced processing, distributed across the human association cortex and implemented in multimodal AI models, is essential to assess the emotional valence of complex social scenes, particularly when it conflicts with simpler image properties.

## Materials and Methods

### Participants

The study includes data from 75 participants. Fifty first-year psychology students (15 males, 17-20 years of age) participated in the behavioral experiment and received course credit. Twenty-two healthy right-handed subjects with normal visual acuity (10 males, 19-47 years of age) took part in the fMRI experiment and received payment for their participation. Three volunteers (1 female, 21-25 years of age), excluding behavioral and fMRI participants, wrote sentences describing each stimulus and received course credit for their help. All participants gave written informed consent. The behavioral experiment was approved by the Social and Societal Ethics Committee of KU Leuven and the fMRI experiment by the Ethics Committee Research of UZ / KU Leuven.

### Stimuli

To devise a proper stimulus set, we first ran a behavioral experiment in which participants rated the valence of the scene, valence of the depicted people, valence of the image as a whole, intensity, and interaction for a set of 148 complex social scenes. For scene valence, participants were asked to annotate the valence of the scene based on the background, objects in the photo, clothing, and attributes that the depicted people hold (everything except body language and facial expression). The valence of people was defined as the facial and body expressions. Finally, for the valence ratings of the image as a whole participants were asked to consider both the scene valence and people valence. These image valence ratings might therefore not be the same as the valence rating assigned to the scene or people in the images. All valence sources were rated from 1 for the most negative to 7 for the most positive. The same definitions as given in the participant instructions were also used as prompts for asking GPT4 to rate the scene/people/image valence for the final set of stimulus images.

To have a balanced stimulus set for studying the human brain, behavior and AI algorithms, we chose a subset of 48 complex social scenes (Fig. 1) that included all possible combinations of the valence of the scene context (positive or negative), the valence of the depicted people/face and body expressions (positive, negative, or ambiguous), and whether there was a direct interaction among those people or not. Positive scene contexts consisted of beach, birth, fair, and wedding and negative scene contexts consisted of funeral, hospital, poverty, and war categories.

We evaluated the reliability of annotations by computing the split-half reliability. We randomly divided participants into two groups, averaged annotations across participants in each group, computed the correlation between them and applied Spearman-Brown correction. Estimated reliability[confidence interval of 95%] for scene, people, and image valence was respectively 0.99[0.98,0.99], 0.99[0.98,0.99], and 0.99[0.98,0.99].

To quantify the incongruence between scene valence and people valence for each stimulus, we defined an incongruence index equal to the absolute difference between corresponding scene valence and people valence. Then, we determined the median of the incongruence index, and labeled stimuli with an incongruence index larger than the median as incongruent stimuli and those with an incongruence index smaller than the median as congruent stimuli (Fig. 1B, gray background).

All images were colorful and cropped to 500 × 700. We equalized the luminance histogram across all the images using the shine toolbox ^40^. In the MRI scanner, these images were shown at a size of approximately 10 degrees of visual angle, on a gray background.

To compare stimuli concerning different low/high-level features, we quantified pair-wise differences between them and computed representational dissimilarity matrices (RDMs). To construct a dissimilarity matrix based on valence (behavioral RDMs), we computed the absolute difference of valences for each stimulus pair. Comparing valence RDMs (Spearman’s correlation between the vectorized off-diagonal elements of RDMs; p values from one-sided permutation tests), we observed larger similarities for congruent stimuli (scene&people: r = 0.79, p < 0.001, scene&image: r = 0.87, p < 0.001, people&image: r = 0.92, p < 0.001) while for incongruent stimuli scene valence was dissociated from people or image valence (Scene&people: r = 0.17, p < 0.05, scene&image: r = −0.06, p > 0.05, people&image: r = 0.70, p < 0.001). To characterize the observed incongruence in stimuli, we built an incongruence RDM; we followed a similar approach as valence RDMs and computed the absolute difference of incongruence indices for each stimulus pair. Comparison of the incongruence RDM with RDMs based on other low/high-level features including scene valence, people valence, image valence, intensity, interaction, color content, pixel intensity, frequency content, Gist model, and scene category did not show significant correlations (Spearman’s correlation and p values from permutation tests, p > 0.05). This finding indicates that all these features were well-controlled between congruent and incongruent images. Full descriptions of the different RDMs and results on their comparisons are provided in Suppl. Note 4.

We also quantified visual features related to the depicted faces and bodies in the images, including the total area of all faces relative to the image area (Face-Area), number of faces (Face-Count), average distance between face pairs relative to the image size (Face-Distance), area of the largest face relative to the image area (Face-Largest), total area of all bodies relative to the image area (Body-Area), number of bodies (Body-Count), area of the largest body relative to the image area (Body-Largest). We used the MTCNN toolbox in Matlab to automatically detect faces, while bodies were segmented and traced manually.

Finally, we asked three volunteers to write sentences describing each stimulus. The instruction was “Can you give a verbal description of this picture that conveys what is shown and happening in the picture?”. We later used this instruction as a prompt to also ask GPT4 to describe the stimulus images.

### Scanning procedures

Data collection consisted of eight experimental runs, three localizer runs, and one anatomical scan, all completed in one long session for each participant (for two subjects, localizers were recorded in a second session).

For the experimental runs, we used a rapid event-related design. Each experimental run included a random sequence of 128 trials: two repetitions of each stimulus image plus 32 fixation trials. Each trial was 3 s. Stimulus trials began with the stimulus presentation for 1500 ms and were followed by 1500 ms of the fixation point. Each experimental run lasted 6 min 44 s. The fixation point was presented continuously at the center of the screen throughout each run. Subjects were instructed to fixate on the fixation point and on each trial, rate the valence of the whole image on a scale of 1-4 with a button press (four buttons with thumb fingers of two hands). The buttons associated with valence ratings were counterbalanced across runs. To make sure that participants can rate the image valence while fixating on the fixation point, they were familiarized with the stimuli before fMRI recording.

For the localizer runs, we used a block design with five stimulus types: body, face, object, scene, and box-scrambled version of the object images. Each run included 20 blocks with four blocks for each stimulus type. The presentation order of the stimulus types was counterbalanced across runs. Each block lasted for 16 s. There was a 10-s blank period at the beginning and end, and three 12-s blank periods between stimulus blocks of each repetition. Each localizer run lasted for 6 min and 16 s. In each stimulus block, there were 18 images of the same stimulus type and two different randomly selected images were repeated. Each image in a block appeared for 400 ms followed by 400 ms fixation. A fixation point was presented continuously at the center of the screen throughout each run. Subjects were instructed to fixate on the fixation point, detect the stimulus one-back repetition, and report it by pressing a response key with their right index finger.

Subjects viewed the visual stimuli through a back-projection screen, and the tasks were presented using MATLAB and Psychtoolbox-3 ^41^.

### Acquisition parameters

Magnetic resonance imaging (MRI) data were collected at the Department of Radiology of the Universitair Ziekenhuis Leuven University Hospitals using a 3T Philips scanner (same imaging parameters as Yargholi & Op de Beeck, 2023), with a 32-channel head coil. Functional images were obtained using a two-dimensional (2D) multiband (MB) T2*-weighted echo planar imaging sequence with an MB of 2, time repetition (TR) of 2 s, time echo (TE) 30 ms, 90◦ flip angle, 46 transverse slices, and a voxel size of 2×2×2 mm3. A high-resolution T1-weighted structural scan was also acquired from each participant using an MPRAGE pulse sequence (1×1×1 mm3 isotropic voxels).

### Preprocessing

fMRIPrep ^43^ was employed for preprocessing anatomical and functional data, using default settings unless otherwise noted. The T1-weighted image was corrected for intensity non-uniformity, skull-stripped, and went through nonlinear volume-based registration to ICBM 152 nonlinear Asymmetrical template version 2009c. Each of the bold runs was motion-corrected, coregistered to the individual’s anatomic scan, and normalized into standard space MNI152NLin2009cAsym. Runs with excessive head motion (frame-wise displacement>2 mm; more than the size of 1 voxel; same threshold as Yargholi & Op de Beeck, 2023 were excluded (2 out of 8 experimental runs for 3 participants; 1 out of 8 experimental runs for 1 participant; and 1 out of 3 localizer runs in two participants). Subjects either did not have frame-wise displacements greater than the predefined threshold or had it repeated multiple times. The rest of the analyses were conducted with SPM12 software (version 6906). As the last step of preprocessing, all functional volumes were smoothed using a Gaussian kernel, 4 mm FWHM. After preprocessing, a run-wise general linear model (GLM) analysis was performed to obtain the beta values for each stimulus image of the experimental and each stimulus type of the localizer runs in each voxel. For the experimental runs, the GLM consisted of the 48 stimulus regressors (boxcar functions at the stimulus onsets with a duration of 1500 ms convolved with a canonical hemodynamic response function), six motion correction parameters (translation and rotation along the x-, y-, and z-axes), frame-wise displacement, cerebrospinal fluid and white matter nuisance signals. For the localizer runs the GLM included the five stimulus regressors (boxcar function at the block onsets with a duration of 16 s convolved with a canonical hemodynamic response function) and the same six motion correction parameters.

### Defining regions of interest (ROIs)

We used two types of ROIs, Human Connectome Project (HCP) ROIs and category-selective networks.

HCP is a whole cortical brain parcellation provided by the Human Connectome Project ^18^, which delineated a total of 360 cortical areas (180 cortical areas per hemisphere). HCP ROIs, cortical parcellation originally defined on the standard cortical surface, were converted to the cortical volume segments using FreeSurfer (version 6.0.1; Fischl, 2012) and then registered to standard space MNI using flirt from FSL (version 5.0.9; Jenkinson et al., 2002). We followed the previously published clustering of these small ROIs into 22 larger ROIs, a clustering that is based on anatomical landmarks, functional similarities or connectivity patterns ^18^. We excluded three ROIs out of these 22, not relevant to the current study (early somatosensory and motor, early auditory, superior parietal). Finally, we considered a division of the remaining 19 ROIs into three large ROIs that broadly characterize three levels of processing of visual information: low-level (primary visual and early visual), mid-level (dorsal stream visual, Ventral stream visual, MT+ complex and neighboring visual areas, paracentral lobular and mid-cingulate, posterior opercular, auditory association, insular and frontal opercular, medial temporal, lateral temporal, temporo-parieto-occipital junction, inferior parietal), and high-level (premotor, posterior cingulate, anterior cingulate and medial prefrontal, orbital and polar frontal, inferior frontal, dorsolateral prefrontal) (Fig. 2B and Fig. S2C). The distinction between mid- and high-level is mostly anatomical as more posterior (mid-level) versus frontal ROIs (high-level), with the exception of posterior cingulate which we lumped with frontal ROIs due to its high intrinsic connectivity with frontal regions as part of the default mode network and its related role in high-level cognitive functions. After analyzing all data with this approach, we verified that this a priori decision of where to categorize posterior cingulate, as well as the decision to omit clusters like superior parietal, have no meaningful impact on the outcome of statistics performed on the three ROIs.

To define category-selective networks, we used contrasts from the localizer runs intersected with functional ^19^ or anatomical (Anatomy Toolbox, Eickhoff et al., 2005) masks in each subject. Details on localizing these category-selective ROIs (employed functional contrasts and masks) are provided in Suppl. Table S4. All lateral and ventral ROIs with the same category-selectivity in both hemispheres were merged to produce large category-selectivity networks. We also used the functional Superior Temporal Sulcus (STS) as a primary social perception ROI^20^, and the anatomical V1 mask to define primary visual cortex as a low-level visual ROI.

All ROIs included at least 25 voxels that surpassed the statistically uncorrected threshold p<0.001 in the relevant functional contrast and were included in the relevant mask (Suppl. Table S4). If the number of surviving voxels was less than 25, a more liberal threshold of p<0.01 or p<0.05 was applied (same procedure as in Yargholi & Op de Beeck, 2023).

### Statistical analysis of the main fMRI experiment

Multivariate and univariate analyses were used to study the representation of scene/people/image valence in the human brain and compared to corresponding representations from behavioral annotations and deep neural networks. Code from https://github.com/KamitaniLab/EmotionVideoNeuralRepresentation supplemented by in-house MATLAB code was used for the following analyses.

### Decoding

We decoded valence (negative vs positive) for scene/people/image adopting the approach of Horikawa et al., 2020. For each valence source, we determined the median of the valence annotations and labeled stimuli with a valence smaller than the median as negative stimuli (−1) and those with a valence larger than the median as positive stimuli (+1). In a leave-one-run-out cross-validation procedure, ridge regression (L2 regularized linear regression) was used to predict stimulus valence from patterns of voxel activity. Ridge regression uses a regularization parameter to constrain the magnitude of the weight coefficients and prevent overfitting. A regularization coefficient was individually estimated for each combination of stimulus labels, ROIs, and subjects; it was optimized based on the model performances obtained by a cross-validation procedure nested within training data using each of 20 possible regularization coefficients (log spaced between 10 and 10000).

Following Horikawa et al., 2020., in each ROI, a set of informative voxels (with a maximum of 500 voxels) showing higher correlations with the valence in the training data were selected to take part in the decoding. Decoding performance was evaluated in the test data (left-out run) by calculating the Pearson correlation between true and predicted labels. This procedure was repeated for each left-out run (6-8 runs), and the correlations were averaged across these repetitions. The resulting correlations were tested for significance using one-sample t-tests across participants, using false-discovery-rate (FDR) correction for multiple comparisons given the number of ROIs ^17^. We obtained decoding performance for all HCP ROIs and investigated the results for 180 ROIs (averaged across left and right hemispheres for HCP ROIs) or the coarser division to 19 ROIs and three large ROIs (averaged across constituting HCP ROIs). In the following, we also used different versions of this decoding approach with variations in defining train and test sets while all other characteristics are the same as mentioned above. We adopted Horikawa et al., 2020 to employ Pearson correlation for evaluating similarity between decoding results and other factors. Other than these cases, Spearman’s correlation was used for comparisons of factors from different modalities.

#### Cross-stimulus decoding, including all stimuli

To control for effects specific to subsets of stimuli, we adopt a cross-stimulus decoding approach. In this approach, if a stimulus (corresponding neural responses and label) is used in the training set, it is not used in the test set. Therefore, in decoding a specific valence source, to maintain a balanced amount of positively and negatively rated valence stimuli, we partitioned the 48 stimuli into six batches, with each batch including four negative and four positive stimuli. Then, for the held-out test run (leave-one-run-out cross-validation), the decoding process was repeated six times; each time just one batch was used as the test set, and the same batch was excluded from training runs. After repeating the process for all batches, we obtained predicted labels for all 48 stimuli (Fig. 2C).

For the comparisons with behavior and AI models, we averaged these predictions across participants in order to gain a more reliable prediction. We refer to these predictions as the group averaged predicted labels. These group-averaged predicted labels were correlated with true labels from behavior and predicted labels from simple and advanced AI models (Fig. 3B and Fig. 6A; significance assessed through permutation tests).

#### Separate cross-stimulus decoding for congruent and incongruent images

We also applied cross-stimulus decoding to congruent and incongruent stimuli separately (Fig. 3A). In these within-congruence cross-stimulus decodings, we faced weakly unbalanced cases (unequal number of stimuli with negative and positive valence) and thus had to exclude a few stimuli to keep the balance. These selections were randomly made for each individual decoding for ROIs and subjects. Then, stimuli were partitioned into batches, each batch including two negative and two positive images.

#### Univariate analysis

We obtained univariate responses of each category-selective network; for each stimulus, beta values were averaged across all runs and all voxels within each network in individual subjects. Then, we computed the correlations between univariate responses and annotated valences (for scene/people/image) (Fig. 4D) or visual features (Face-Area, Face-Count, Face-Distance, Face-Largest, Body-Area, Body-Count, Body-Largest) (Fig. 4B), applied t-tests across participants, and corrected p values for multiple comparisons across networks (FDR) ^17^.

#### Computational simulation with the AI models

To understand the representation of our stimuli in AI models, we tried different approaches and compared those with behavioral or neural representations. We examined various AI models, simple ones like AlexNet ^22^ and EmoNet ^4^, more advanced models such as CLIP ^23^, and finally Large Language Models like Llama ^24^ and GPT4 ^25^.

#### Valence predictions by AI models

In this section, we explicitly and directly queried and compared the valence annotations in AI models. As a first model for predicting the valence of each image, we used the EmoNet model ^4^. They retrained the weights in the final fully connected layers of AlexNet, changed its objective from recognizing object classes to identifying emotion categories, and called the new model EmoNet. Then, they constructed a predictive model of image valence from the last fully connected layer of EmoNet using partial least square regression. As a second and more advanced AI model, we used CLIP and augmented it with a single layer at the output to provide us with the prediction of image valence. The CLIP models were obtained by training five models per starting learning rate lr_0_∈ [10^−3^, 10^−4^], using Cross Entropy Loss and Adam optimizer with default parameters and the custom learning rate update rule lr_e_ = lr_0_/ (e//3)+1 with lr_e_ the learning rate at epoch e. The data used was the publicly available part of the FindingEmo dataset, randomly split using an 80/20 train/test split ratio ^47^. Out of all the trained models, we selected the one with the best Mean Absolute Error on the test set as the winner. The training or test images of the FindingEmo dataset do not include any of the images used in the current study. To have a less confounded comparison between Emonet and CLIP, we also augmented EmoNet as provided by Kragel et al., 2019 with a single layer at the output and finetuned it on FindingEmo using the same approach as for CLIP (see Supl. note 8 for resluts). We also used Llama and GPT4 as a large language model and asked for scene/people/image valence ratings for each image one-by-one, providing the same instructions (different prompts for different sources of valence) and definitions for valences used in human annotations. To make sure that each image is annotated independently, We did not have memory over consecutive requests. We employed Llama3.2-11B-Vision-Instruct from (https://huggingface.co/models). The annotations of GPT4 were collected in January 2024 using the updated “gpt-4-0613” model (https://platform.openai.com/docs/models). For 20% of annotations, we repeated ratings employing chat GPT4 and obtained results were the same as when we used GPT4 API. GPT4 refused to rate valences for 8 images presumably due to content moderation (ethical and safety concerns).

Then, we computed the correlation between valence predictions from AI models and behavioral annotations for scene/people/image valence, using Spearman’s correlation. The significance of these correlations was verified by using permutation tests. The procedure was repeated for (in)congruent stimuli separately (Fig. 5).

We also compared the AI-predicted and behaviorally annotated valences with neurally decoded scene/people/image valence from cross-stimulus decoding in low/mid/high-level ROIs for (in)congruent stimuli through Spearman’s correlation. For this comparison, we used the group averaged predicted valence obtained from cross-stimulus decoding The significance of these correlations was verified by using permutation tests (Fig. 6).

## Acknowledgment

The authors would like to thank Yannik Timar and Deniz Erdil for some preliminary analysis and Aicha Boutachkourt for her help with the fMRI recordings. This work was supported by Fonds voor Wetenschappelijk Onderzoek 12A6122N, G0D3322N, and G073122N; the KU Leuven research council (interdisciplinary project IDN/21/010); and long-term structural Methusalem funding by the Flemish Government (METH/24/003).

## Supplementary

### Supplementary Notes 1. Links to stimulus images

Table S1. Includes links to all 48 stimulus images.

**Table S1.**
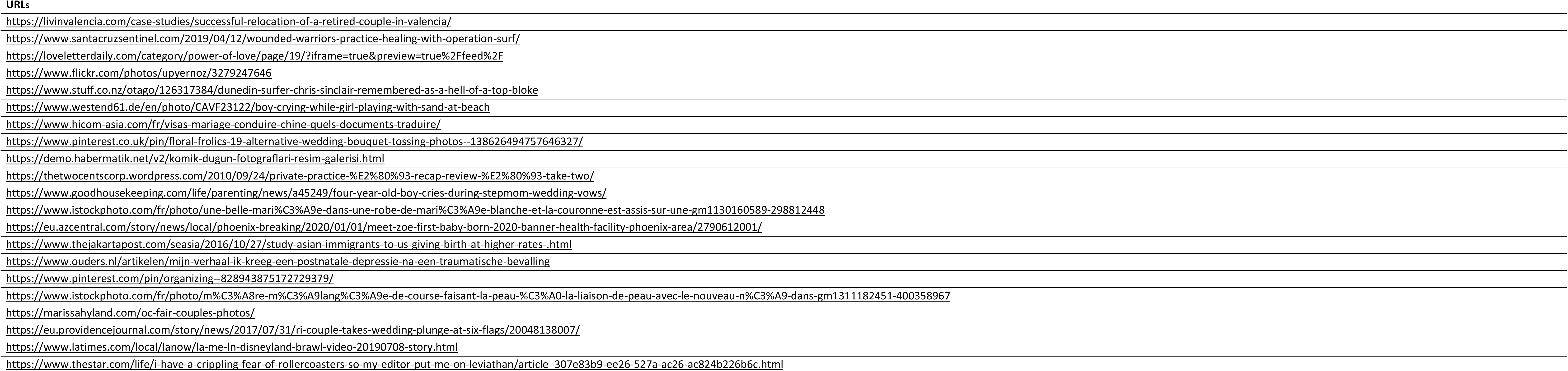
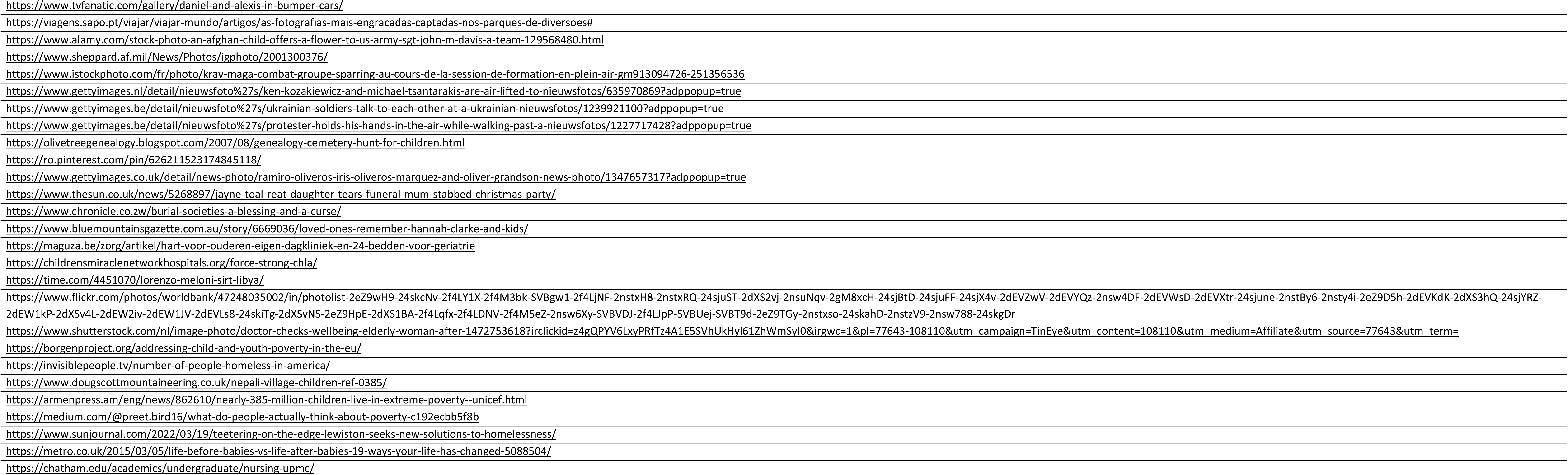
Links to stimulus images.

### Supplementary Notes 2. Cross stimulus decoding in 19 cortical regions

To further pinpoint the location of the strongest decoding results, we applied a previously proposed clustering of HCP ROIs (Glasser et al., 2016) in 19 regions covering all cortex except somatosensory and motor, early auditory, and superior parietal. One-sample one-sided t-tests corrected for multiple comparisons across ROIs (FDR) (Benjamini & Hochberg, 1995) were applied to averaged decoding performances in these regions. For scene valence, ventral stream visual, insular and frontal opercular, temporo-parieto-occipital junction, inferior parietal, posterior cingulate and for people valence, posterior cingulate and dorsolateral prefrontal ROIs showed significant correlations between true and predicted valences. Therefore, posterior cingulate significantly decoded both scene and people valence, the other significant prediction for people valence was derived in a more anterior ROI in comparison to scene valence. Scene valence was significantly predicted in more posterior ROIs.

**Fig. S1.**
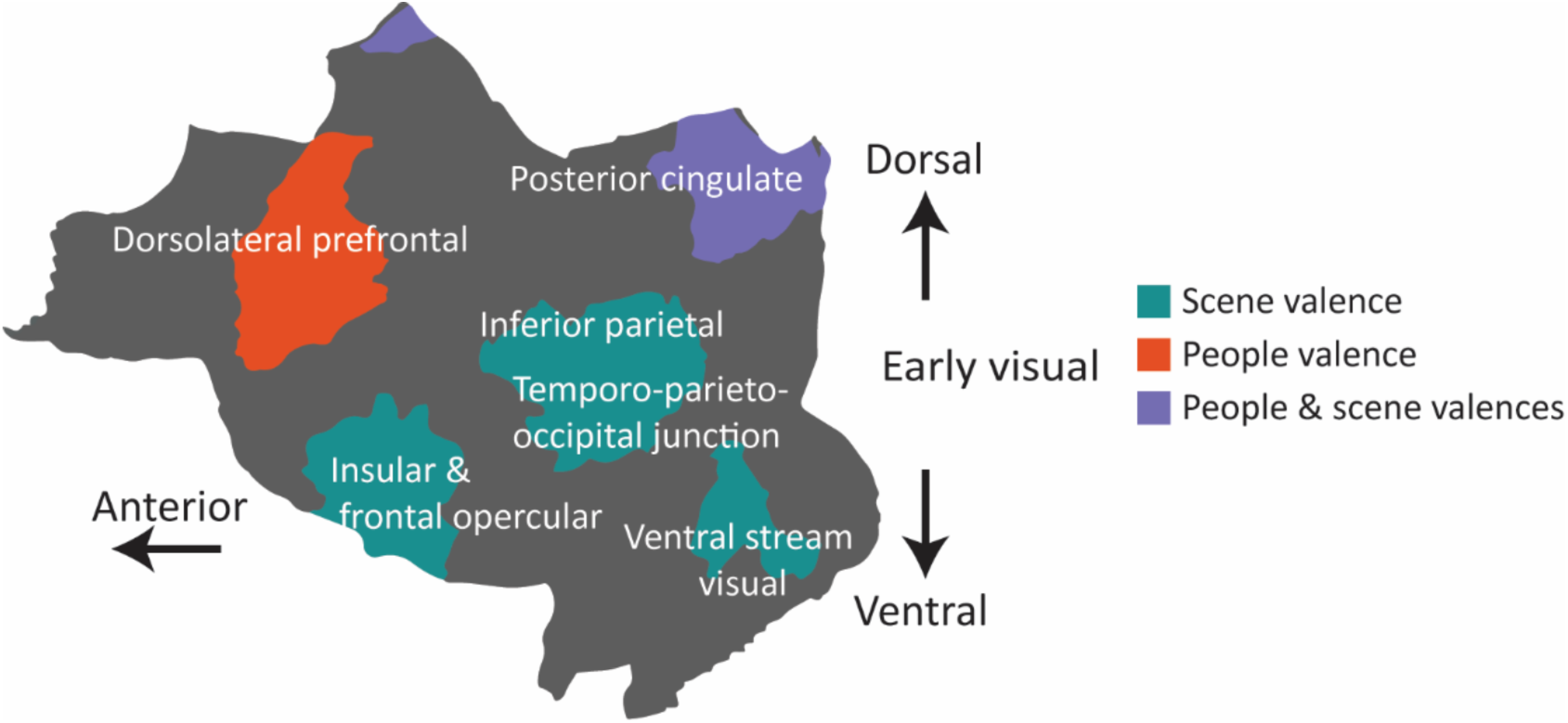
Regions with significant cross-stimulus decoding. These ROIs showed significant accuracy (Pearson correlation between true and predicted valences) in cross-stimulus decoding for scene/people valence (one-sample one-sided t-tests, FDR corrected, p < 0.05).

### Supplementary Note 3. Simple decoding

To evaluate the representation of scene/people/image valence in different brain regions, we employed a decoding approach. Using ridge regression, we fitted the decoding models to the fMRI data acquired while subjects viewed the stimulus images and rated the valence of the whole image. We predicted positive vs. negative valences for scene/people/image in HCP (Human Connectome Project) regions following a leave-one-run-out approach (Fig. S2A). Decoding performance was evaluated by calculating the Pearson correlation between true and predicted valences ^16^ and tested for significance using one-sample one-sided t-tests corrected for multiple comparisons across ROIs (FDR) ^17^. Fig. S2B shows the decoding performance for 180 HCP ROIs (averaged across left and right hemispheres) on the flattened cortical surface. Many regions are decoding the valences significantly above chance. Predictions for all valences are more accurate in low-level regions and scene valence is the most accurately decoded valence. Note that the higher decoding in low-level regions is not due to confounds between differences in valence and differences in low-level properties as captured, e.g., by GIST (Suppl. Fig. S3).

To summarize these results, we considered a division of HCP ROIs into three large ROIs: (i) low-level visual regions (retinotopic areas V1-V4), (ii) mid-level association areas mostly in posterior cortex, and (iii) high-level ROIs involved in control processes and mostly in anterior cortex. We averaged decoding performances across their constituting HCP ROIs (Fig. S2C). These results show a significant correlation between true and predicted labels in all regions for all valences, with higher correlations for low-level ROIs (primary visual and early visual) and scene valence. The variation of decoding across sources of valence and ROIs was tested statistically in a 2-way ANOVA with the source of Valence (scene/people/image) and Region (low/mid/high-level) as within-subject factors. We found a significant main effect of Valence (F(2, 84) = 25.06, p = 6.88e-08), region (F(2, 84) = 81.34, p = 3.55e-15), and Valence*Region interaction (F(4, 84) = 28.06, p = 8.33e-15).

**Fig S2.**
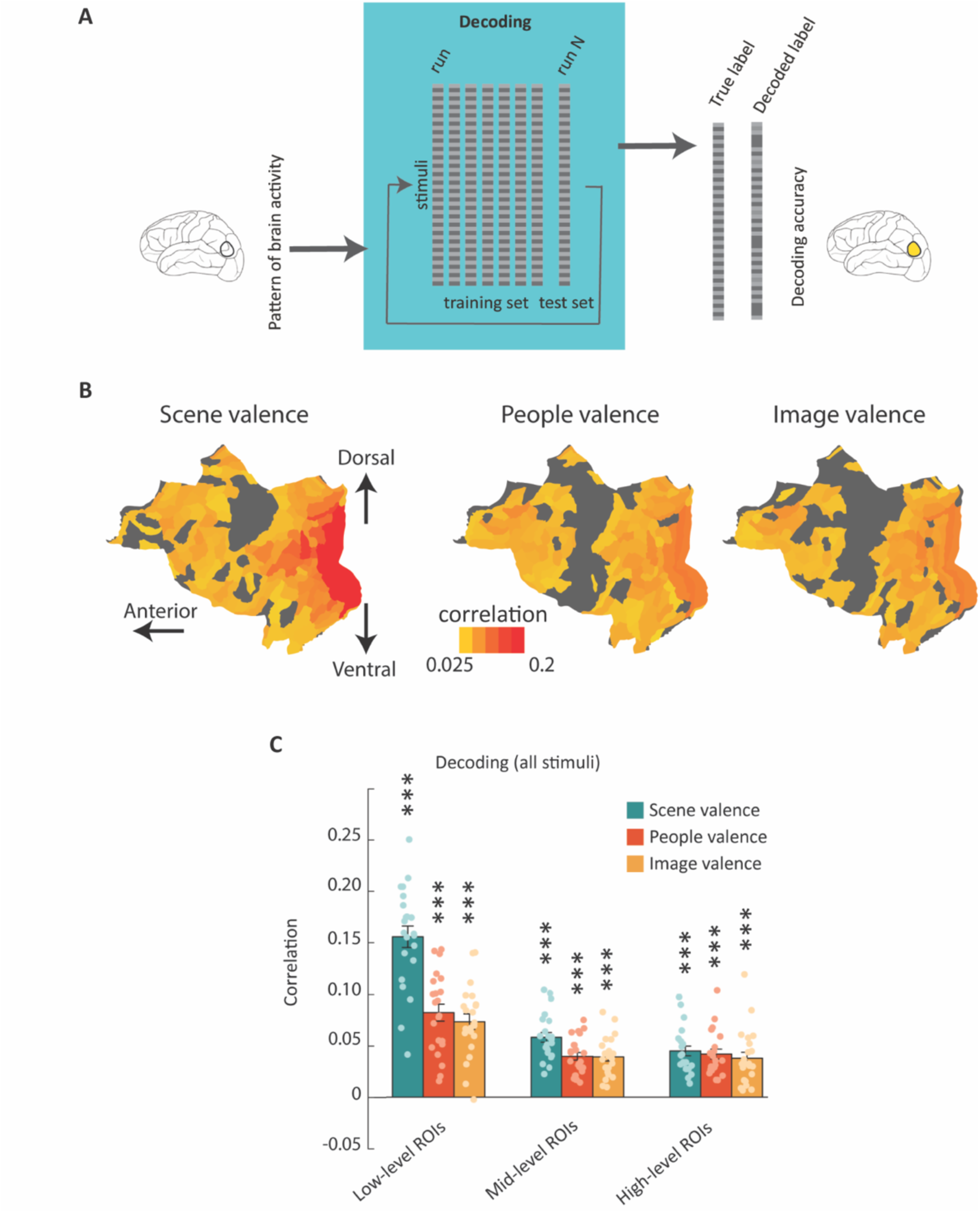
Simple decoding approach and performance. A, Schematic presentation of simple decoding approach. Decoders predicted positive vs. negative valences following a leave-one-run-out approach. B, Simple decoding performance in HCP ROIs. Colorful ROIs had significant decoding accuracy for scene/people/image valence (Pearson correlation between true and predicted valences, one-sample, one-sided t-tests, FDR corrected, p<0.05) C, Performance of simple decoding for scene/people/image valence in low/mid/high-level ROIs. The bar plots show the mean decoding accuracy (Pearson correlation between true and predicted valences). Dots represent individual subjects and error bars indicate SEM (* p < 0.05, ** p < 0.01, *** p < 0.001, one-sample one-sided t-test, FDR-corrected).

### Supplementary Note 4. Comparison of behavioral and model RDMs

To compare stimuli concerning different low/high-level features, we quantified pair-wise differences between them and computed representational dissimilarity matrices (RDMs). The off-diagonal of the RDMs was vectorized, and Spearman’s correlation between dissimilarity vectors was then calculated and tested (p values from one-sided permutation tests). Fig. S3 shows the correlation between vectorized RDMs (colorful cells remark significant correlations survived one-sample one-sided permutation tests, p<0.05).

For constructing a dissimilarity matrix based on behavioral annotations including scene/people/image valences, intensity, and interaction and incongruence index (behavioral RDMs), we computed the absolute difference of the corresponding values for each stimulus pair. To compare stimulus images based on their color content, we obtained histograms for red, green, and blue colors, applied pairwise Euclidean distance to compute RDM for each color and averaged these three RDMs to have a final color RDM.

As a measure of low-level shape properties, pixel-wise dissimilarity was investigated (OP de Beeck et al., 2008). We vectorized the pixel intensity matrix of each stimulus image and calculated the pairwise Euclidean distance to obtain an RDM based on pixel intensity.

The frequency content of stimulus images was also examined in a similar approach to pixel intensity. In this case, the primary feature vector was the vectorized version of the amplitude matrix produced by the Fourier transform.

We also employed the GIST model which effectively captures scene statistics and accurately represents responses in lower visual areas (Groen et al., 2013; Oliva & Torralba, 2001). Correlation distance between GIST features was used to build the GIST RDM.

Category RDM was based on the eight scene categories in the stimulus set (beach, birth, fair, wedding, funeral, hospital, poverty and war). RDM values were one (+1), except for stimuli of the same category which had RDM values equal to zero (0).

**Fig S3.**
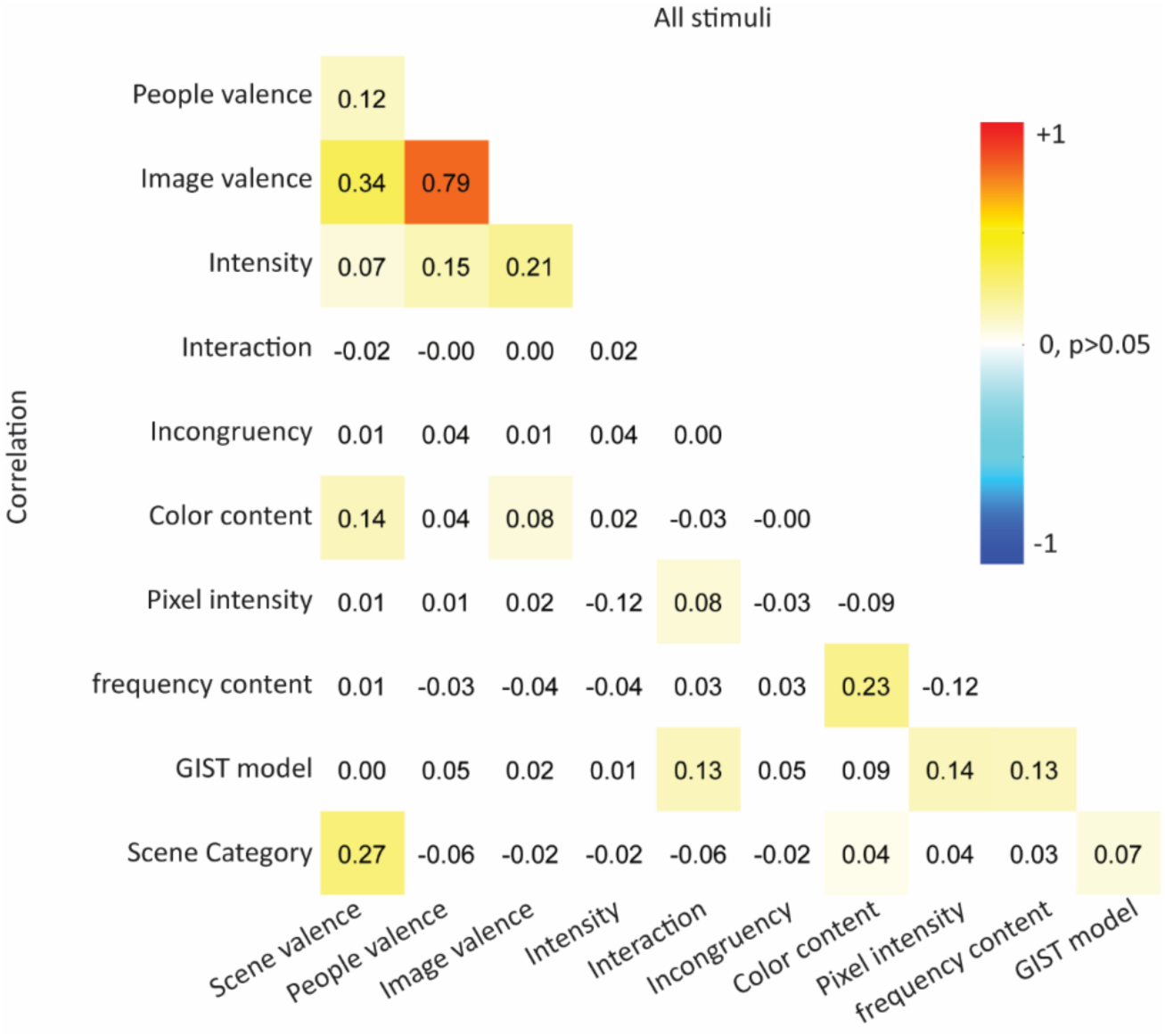
Correlation between behavioral and model RDMs. Spearman’s correlation between RDMs (colorful cells remark significant correlations survived one-sample one-sided permutation tests, p<0.05).

### Supplementary Note 5. cross-stimulus decoding in category-selective networks

we also employed cross-stimulus decoding to predict positive vs negative valences for scene/people/image in these regions. Results did not reveal any significant cross-stimulus decoding, except for scene valence in the scene network (Suppl. Fig. S4).

**Fig. S4.**
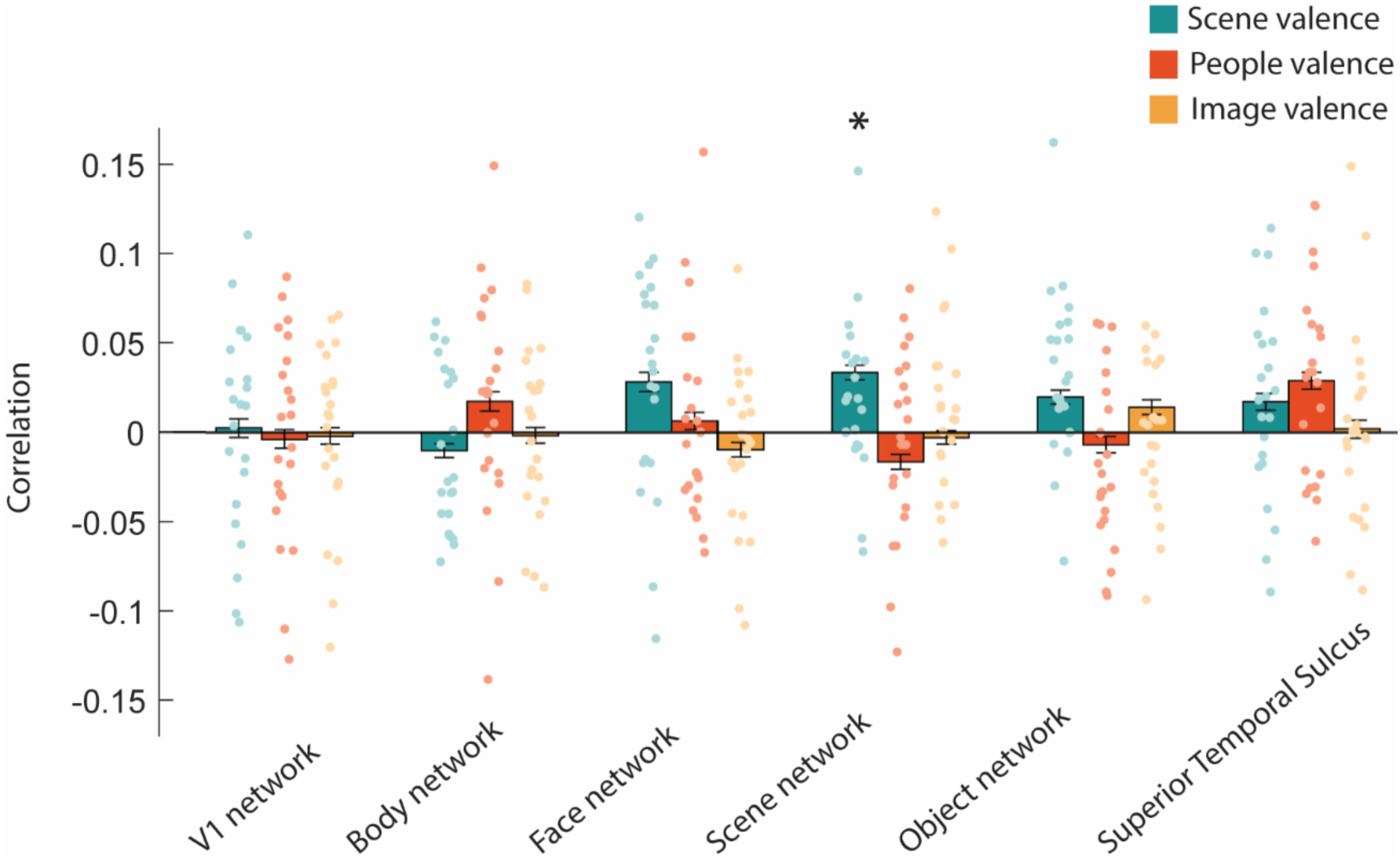
Performance of cross-stimulus decoding in category-selective networks. The bar plot shows the mean decoding accuracy (Pearson correlation between true and predicted valences) in brain networks for scene/people/image valences. Dots represent individual subjects and error bars indicate SEM (* p < 0.05, one-sample one-sided t-test, FDR-corrected).

### Supplementary Note 6. Representational similarity analysis

#### Neural representation of valence in low/mid/high-level ROIs

To examine the neural representation of scene/people/image valence in different ROIs, we compared neural dissimilarity in the multivoxel patterns elicited by individual images with predictions from behavioral annotations i.e. the representational dissimilarity matrices (RDMs) based on neural activity in different ROIs with behavioral RDMs based on scene/people/image valence (same procedure as in Yargholi & Op de Beeck, 2023). Neural RDM for each ROI included the pairwise Mahalanobis distance between activity patterns (b weights) of the ROI for different stimuli ^51,52^. The off-diagonal of the neural and behavioral RDMs was vectorized, and Spearman’s correlation between dissimilarity vectors was then calculated and compared. We tested the significance of correlation values between neural and behavioral RDMs across subjects using one-sample t-tests. Then, p values were corrected for multiple comparisons across all ROIs (false discovery rate [FDR]) ^17^. The result is shown in Fig. S5A. There are significant positive correlations only with scene valence in low-level ROIs (p < 0.001) and mid-level ROIs (p < 0.01).

#### Representations in the last hidden layers of AlexNet, EmoNet, and CLIP only capture scene valence, to a limited degree

For open-source models (AlexNet, EmoNet, and CLIP), we obtained representations of stimulus images in the last hidden layers. Then to compare these representations with each other and also with behavioral annotations, we employed representational (dis)similarity analysis (RSA) and used model representational dissimilarity matrices (RDMs) and behavioral RDMs. To build model RDMs, we computed the correlation distance between the activations of the units in the last hidden layers for each pair of stimuli. For constructing behavioral RDMs we computed the absolute differences of valences for each stimulus pair.

The off-diagonal of the RDMs was vectorized, and Spearman’s correlation between dissimilarity vectors was then calculated and compared. We tested the significance of correlation values between RDMs using permutation tests. The results are shown in Fig. S5B relevant first finding concerns the correlations among the three behavioral RDMs, for Scene, People, and Image valence (top two rows). The behavioral RDM for image valence is correlated with both scene valence and people valence, with a stronger correlation for people valence. People and scene valence are almost fully dissociated (0.12), which was an explicit aim in the construction of the stimulus set. In terms of correlations between the model RDMs and the behavioral RDMs, we observe two effects: (i) Correlations with scene valence are higher than correlations with the other sources of valence; and (ii) Correlations are higher for CLIP than for the other two models. Thus, EmoNet, which is the previously developed benchmark model for assessing image valence, is outperformed by CLIP which is characterized by a training with images and their verbal labels. Yet, even CLIP mostly captures scene valence. Regarding correlations among the models, AlexNet and EmoNet RDMs are correlated at a level of 0.46, higher than the correlations of these two models with CLIP. Most importantly, the final hidden layers of these models do not contain a representation that captures the image valence of social scene images in an obvious way.

#### Representations in the verbal descriptions provided by humans and multimodal AI models

We employed verbal descriptions to access the representational space of complex social scenes in human and AI models. We asked/prompted volunteers and GPT4 to provide written descriptions for stimulus images and also obtained corresponding CLIP descriptions (https://github.com/openai/CLIP.git). Then, we used Google Universal Sentence Encoder (GUSE) to map sentences to GUSE’s latent space, a semantic space ^53^, averaged latent vectors across participants, and computed the correlation distance to build RDMs based on latent vectors (GUSE-Human, GUSE-CLIP, GUSE-GPT4). We compared these GUSE RDMs with each other and with behavioral RDMs (as discussed in the previous section) using Spearman’s correlation, with significance verified using permutation tests. As Fig. S5C shows all GUSE RDMs had larger correlations with scene valence than people/image valence and GUSE-GPT4 is more similar to GUSE-Human than GUSE-CLIP. Yet, these findings from using verbal descriptions might not be a strong indication that the AI models fail to capture image valence, as the semantic embeddings of human verbal descriptions fail so too. We might need a more direct way to extract valence representations.

**Fig. S5.**
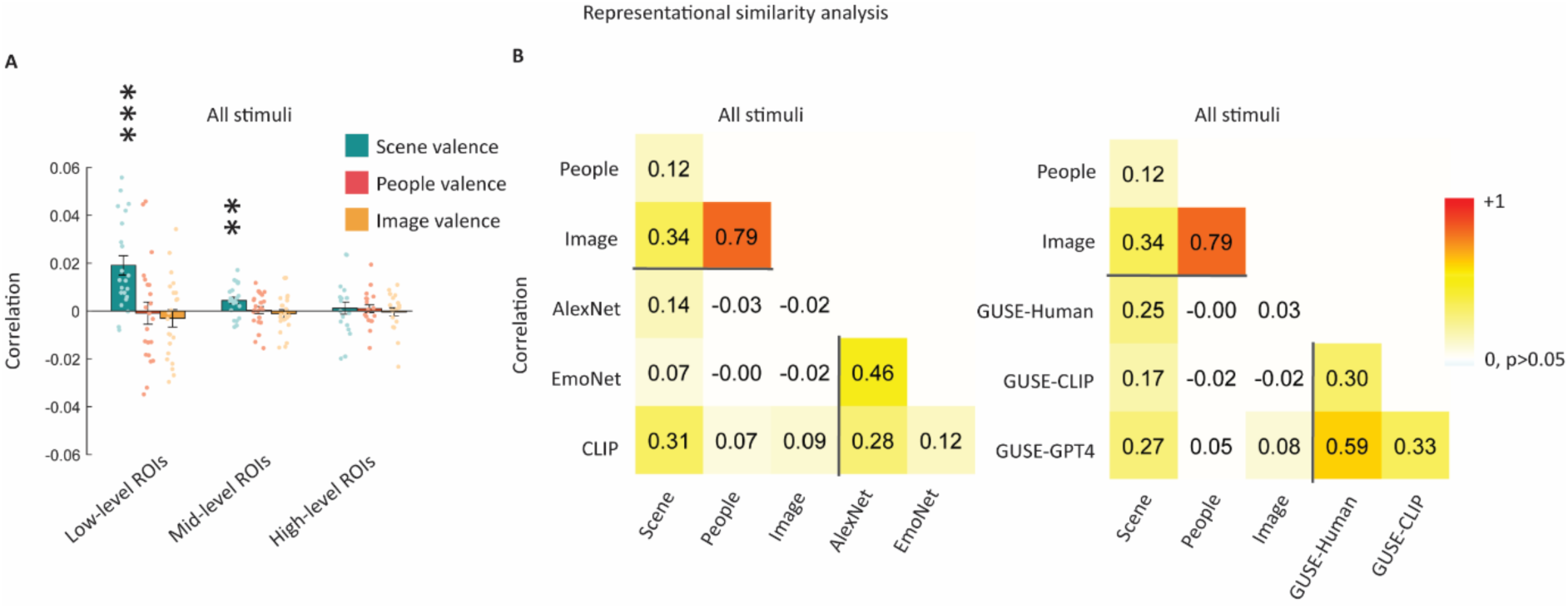
Representational similarity analysis between behavioral RDMs and neural RDMs or model RDMs. A, Representational similarity and effects of scene/people/image valence in the low/mid/high-level ROIs for all stimuli. The bar plot represents the mean Spearman’s correlations between neural RDMs for individual subjects (dots) and behavioral RDMs. Error bars indicate SEM. ***p,0.001; *p,0.01; *p,0.05; one-sided one-sample or two-sided paired t test. B, Representational similarity between behavioral RDMs and model RDMs (representations in the last hidden layers). Correlation matrix for the dissimilarity vectors (colorful cells p < 0.05, permutation test). C, Representational similarity between behavioral RDMs and model RDMs (representations in the verbal description). Correlation matrix for the dissimilarity vectors (colorful cells p < 0.05, permutation test). Values above the horizontal black line are correlations merely among behavioral RDMs and values on the right of the vertical black line are correlations merely among model RDMs.

### Supplementary Note 7. Functional contrasts and masks used to localize category-selective networks

**Table S4.** Functional contrasts and masks used to localize category-selective networks.

| ROIs | Contrast | Mask |
| --- | --- | --- |
| Body network |  |  |
| IEBA & rEBA | body>(face+object) | functional atlas |
| Face network |  |  |
| IFFA & rFFA | face>(body+object) | functional atlas |
| IOFA & rOFA | face>(body+object) | functional atlas |
| ISTS & rSTS | face>(body+object) | functional atlas |
| Object network |  |  |
| lLOC & rLOC | object>scramble | functional atlas |
| Scene network |  |  |
| IPPA & rPPA | Scene>object | functional atlas |
| IRSC & rRSC | Scene>object | functional atlas |
| IOPA & rOPA | Scene>object | functional atlas |
| Superior Temporal Sulcus |  |  |
| ISTS & rSTS (Deen et al., 2015) | All conditions>Fixation or All conditions<Fixation | functional atlas |
| EVC |  |  |
| IV1 & rV1 | All conditions>Fixation | anatomical atlas |

EBA (extrastriate body area), EVC (early visual cortex), FFA (fusiform face area), LOC (lateral occipital complex), OFA (occipital face area), OPA (occipital place area), PPA (parahippocampal place area), RSC (retrosplenial complex), STS (superior temporal sulcus)

### Supplementary note 8. Comparison between CLIP and EmoNet

The comparison between the successful CLIP-Image and the unsuccessful EmoNet-Image involves multiple confounding differences. Some of these differences are inherent to these models and present in all comparisons in the literature between CLIP and AlexNet (the latter is the backbone on which EmoNet was developed), including differences in architecture and initial training (e.g., multimodal input in the case of CLIP). However, we added one new difference. EmoNet-Image was developed from AlexNet by building a model for emotion prediction based upon the last layer, using a training set consisting of relatively simple natural images. In contrast, we developed CLIP-Image from CLIP by augmenting the model with an additional layer trained on valence labels of complex social scene images as available in the recently developed FindingEmo dataset ^47^. Possibly, this training on complex social scenes would be responsible for the success of CLIP-Image, compared to the training on much simpler images in the case of EmoNet-Image. Would EmoNet-Image capture image valence for incongruent images when trained on FindingEmo? To test this hypothesis, we augmented EmoNet as provided by Kragel et al., 2019 with a single layer at the output and finetuned it on FindingEmo using the same approach as for CLIP-Image. However, for incongruent stimuli, the predicted image valence of this network was only significantly correlated with the original EmoNet-Image (permutation test, p < 0.05), and not with behaviorally annotated image valence. The limited capability of EmoNet to capture the valence of complex social scenes could be due to the architecture or its original training protocol, both of which lack the multimodal aspect that is inherent to CLIP.

